# The IRE1α-endonuclease regulates PD-1 expression through a novel XBP1/miRNA-34a axis within Natural Killer cells

**DOI:** 10.1101/2023.02.26.530063

**Authors:** Karolina Bednarska, Gayathri Thillaiyampalam, Sally Mujaj, Jamie Nourse, Jay Gunawardana, Muhammed B. Sabdia, Qingyan Cui, Lilia M. de Long, Frank Vari, Maher K. Gandhi, Alexandre S. Cristino

## Abstract

Activation of the IRE1α-endonuclease is critical for Natural Killer (NK)-cell function. We describe a novel regulatory role for IRE1α-endonuclease in fine-tuning NK-cell effector functions through an inter-connected activation of the transcription factor XBP1s and inhibition of microRNA-34a-5p (miR-34a-5p) to modulate PD-1 immune checkpoint expression. NK-cells, when exposed to cancer cells, activate IRE1α-endonuclease mediated decay of miR-34a-5p. This reduces miR-34a-5p and consequently increases the expression of the target genes XBP1 and PD-1. IRE1α-endonuclease activation not only enhances NK-cell effector function but also promotes PD-1 expression. PD-1 is itself directly regulated by miR-34a-5p, which binds to the 3’UTR of PD-1 messenger RNA to repress PD-1 protein at the NK-cell surface. IRE1α-pathway activation is impaired in the NK-cells of patients with Hodgkin Lymphoma, and miR-34a-5p and PD-1 expression are inversely correlated. The IRE1α-pathway plays a dual role in regulating the XBP1/miRNA-34a axis and PD-1 expression within NK-cells, that is disrupted in cancer patients.

## 1 Introduction

Inositol-requiring protein 1α (IRE1α) is a serine/threonine-protein kinase/endoribonuclease stress sensor that operates within the unfolded protein response system. Under endoplasmic reticulum (ER) stress, IRE1α undergoes oligomerisation and subsequent autophosphorylation that triggers its endonuclease activity. The latter enables processing of its principal substrate, X-Box Binding Protein 1 (*XBP1*) mRNA, via excision of a 26-nucleotide intron and generation of the active form, the transcription factor *XBP1*-spliced (*XBP1s*) (1). The IRE1α-pathway is a highly evolutionarily conserved cell signaling pathway with multiple roles in both innate and adaptive immune cells including plasma cells, dendritic cells, eosinophils, macrophages, T-cells and NK-cells (2–9). Within these immune cell subsets, the functional role of the IRE1α-pathway is highly cell context-specific.

Additionally, it has been established in cell-free systems that the IRE1α-endonuclease recognizes a consensus sequence (CUGCAG) motif, that enables it to cleave selected microRNAs (miRNAs) during ER stress (i.e., miR-17, miR-34a, miR-96 and miR-125b) (10, 11). MiRNAs function in part by targeting the 3’ untranslated regions (3’UTRs) of messenger RNAs (mRNAs) for degradation and/or translational repression to ‘fine-tune’ expression of various proteins. Of particular interest within an immune context, is miR-34a-5p which binds to the 3’UTR of PD-L1 in the malignant cells of various solid malignancies and blood cancers (12–14). The regulation of the miR-34a-5p/PD-L1 axis by diverse stimuli such as the p53 pathway and Epstein-Barr virus within a range of malignant cell types strongly implicates it as an important regulatory hub that can be manipulated for therapeutic benefit (12–15).

Interestingly, miR-34a-5p is also expressed by a range of immune effector cells including dendritic cells, macrophages, mast cells, B and T-cells (16–19). Importantly, miR-34a-5p is also a key regulator of multiple inhibitory/activating receptors and chemokines in NK-cells (20). Recently it has been shown that the IRE1α-pathway mediates NK-cell responses towards viral infections and murine melanoma tumor models *in-vivo*, as well as cytotoxic function and proliferation of human NK-cells from healthy blood donors (3, 9). To our knowledge, there is minimal data on the functional role of the IRE1α-pathway in primary NK-cells from patients with human malignancies, and the role of a potential IRE1α-miR-34a-5p axis in NK-cells remains unexplored.

NK-cells are an important subset of innate immune effector cells that possess the unique ability to lyse malignant cells without previous sensitization (21, 22). Major histocompatibility complex (MHC) class I molecules, frequently observed to be downregulated or missing in malignancies such as Hodgkin Lymphoma (HL), are associated with enhanced NK-cell killing (23). Whereas only a small fraction of intratumoral T-cells will express the relevant T-cell receptor capable of recognizing a malignant cell in an antigen-specific manner, the majority of NK-cells are able to exert anti-tumoral cytotoxicity. Furthermore, their cytotoxic capabilities allow them to kill tumor cells even at relatively low ratios (24). For these reasons, strategies that may enhance endogenous NK-cell immunity such as PD-1 immune checkpoint blockade, have recently attracted attention for their potential use as immune-based therapies (25, 26).

However, there are contradictory reports of PD-1 expression on human NK-cells. Its surface expression is detected on NK-cells in multiple solid organ and blood cancers (25–31). Conversely, minimal surface expression of PD-1 on both peripheral and intratumoral NK-cells of several tested cancer types has also been reported (32). Several studies may help explain these contradictory findings. A comprehensive analysis of both mRNA and protein PD-1 expression in either tumor-associated or healthy donor-derived NK-cells showed different PD-1 isoforms are present in NK-cell cytoplasm, indicating that NK-cells might be able to swiftly externalize PD-1 to the surface following appropriate stimuli (33). Furthermore, it was recently shown that NK-cells can acquire functional PD-1 transferred from cancer cells via trogocytosis (34). Although the transcriptional mechanisms that regulate PD-1 expression in NK-cells are also of great importance and interest, they are not yet well understood. In this study, we sought to investigate the role of IRE1α-pathway underpinning NK-cell effector functions and its mechanistic basis to regulate immune checkpoint PD-1 expression.

## 2 Results

### 2.1 Activation of the IRE1α-endonuclease in NK-cells reduces levels of miR-34a-5p

To confirm that the IRE1α-pathway can be activated in NK-cells in response to blood cancer cell-lines, we co-incubated the NK-cell-line SNK10 with the Hodgkin-Reed Sternberg (HRS) cell-lines HDLM2 (MHC-I deficient) or KMH2 (MHC-I intact), and the NK-cell sensitive target leukemia cell-line K562 (MHC-I and II deficient). Splicing of the transcription factor XBP1 to its active form XBP1s was used as the functional read-out to indicate IRE1α-endonuclease activation. The mRNA expression of XBP1s in K562, KMH2 or HDML2 activated SNK10 cells increased by at least 2-fold in comparison to unstimulated cells (**Fig. 1A**). Similarly, significant increase in XBP1s protein expression was observed in stimulated SNK10 cells (**Fig. 1B; Fig. S1A**). However, the increase of XBP1s protein expression in SNK10 cells stimulated with both HL cell-lines (KMH2 and HDLM2) was far more pronounced (up to ~5-6-fold) than with K562 leukemia cells (~2-fold; **Fig. 1B; Fig. S1A**).

**Figure 1:**
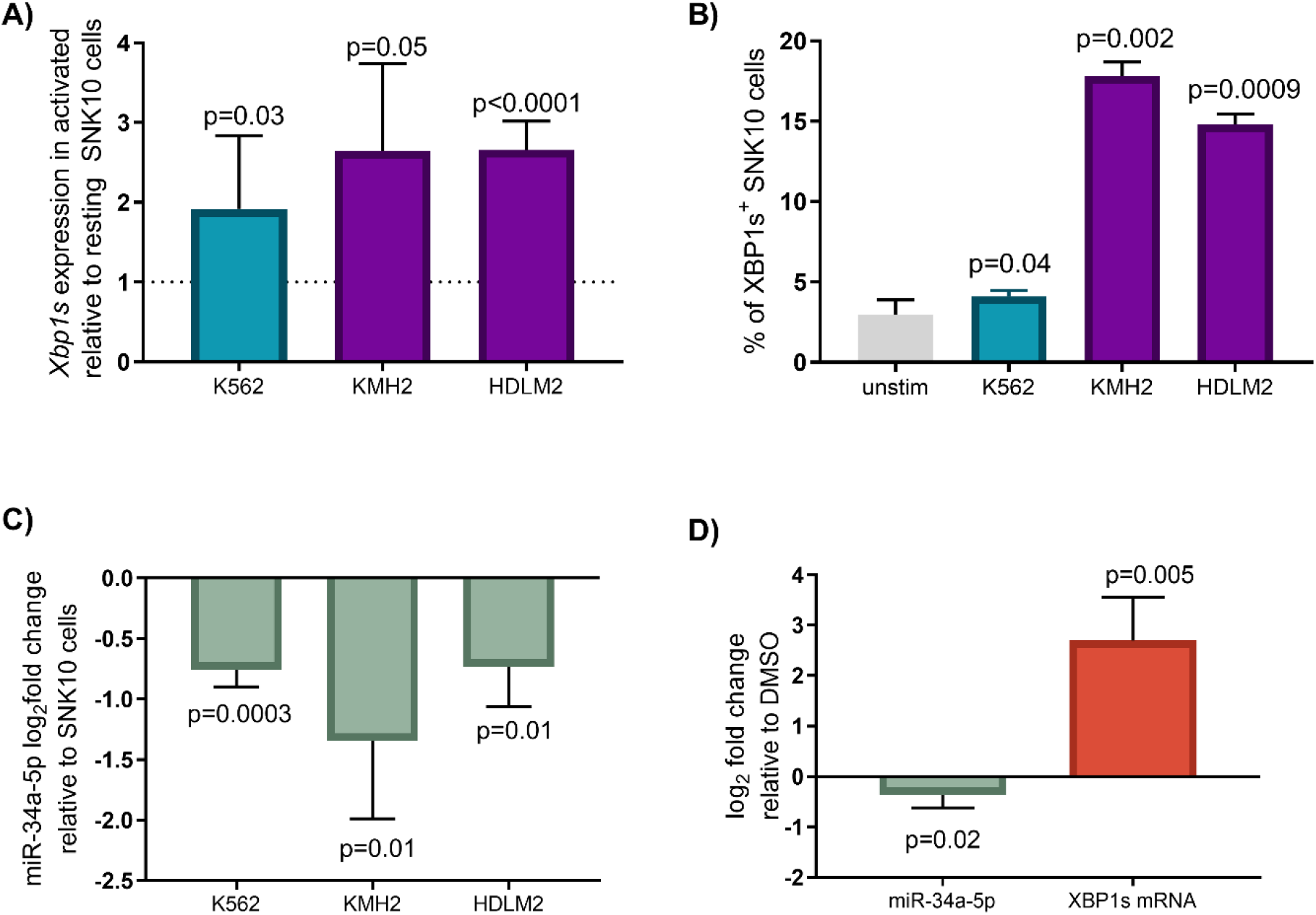
IRE1α-pathway downregulates miR-34a-5p in NK-cells following activation by Hodgkin Reed-Sternberg and leukemia cells. **(A)** IRE1α-pathway activation was confirmed by qRT-PCR analysis of the XBP1 spliced isoform (XBP1s) upon SNK10 NK-cells co-incubation for at least 2 hours with leukemia (K562) or Hodgkin Lymphoma (HDLM2 and KMH2). The bars show changes in the expression of XBP1s in SNK10 cells activated by target cells, relative to resting SNK10 cells (shown by dotted line; B2M is used as endogenous control). **(B)** The expression of XBP1s protein is also upregulated in NK-cells stimulated with different target cells (relative to unstimulated NK-cells). Graph shows the mean ± SEM FACS data of % XBP1s^+^ SNK10 cells. **(C)** qRT-PCR analysis confirms that miR-34a-5p expression is down-regulated in SNK10 cells stimulated with all three cancer cell-lines (RNAU6 used as endogenous control and SNK10 unstimulated as reference condition). A one tailed paired t-test was used for statistical analysis. **(D)** HEK293T cells treated with the ER stressor thapsigargin (TG), to activate IRE1α-pathway, significantly increased XBP1s mRNA while supressing miR-34a-5p expression showing this mechanism is conserved beyond immune cells. Unpaired t-test two tailed used for statistical analysis.

In addition to activating XBP1s splicing, it has been shown in cell-free systems that the IRE1α-pathway also plays a role in the degradation by cleavage of a subset of CUGCAG motif-containing miRNAs including miR-34a (11). Next, we sought to establish if degradation of miR-34a is operative within NK-cells. Firstly, we measured miR-34a-5p expression by qRT-PCR in SNK10 cells stimulated with these three different target cell-lines: K562, KMH2 and HDLM2. This showed significant downregulation of miR-34a-5p in SNK10 cells after stimulation with all three targets cell-lines (**Fig. 1C**). To confirm the mechanism of IRE1α-mediated regulation of XBP1s and miR-34a-5p is conserved in cells other than NK-cells, we tested IRE1α activation in HEK293T cells using a well-known ER stress inducer Thapsigargin (TG). Activation of the IRE1α-pathway in HEK293T cells by TG down-regulated miR-34a-5p expression, whilst XBP1s isoform was upregulated (**Fig. 1D**). These results strongly suggest that IRE1α endonuclease activity plays a dual role in the control of expression of XBP1s and miR-34a-5p within NK-cells as well as other cell types.

### 2.2 miR-34a-5p targets the 3’UTRs of XBP1 and PD-1 mRNAs to suppress their expression

Next, we performed a computational analysis to identify putative target sites for all four miRNAs known to be regulated by IRE1α (miR-17, miR-34a, miR-96 and miR-125b) (11), in the 3’UTR region of XBP1 and PD-1 mRNAs. Adopting a stringent criterion for miRNA target site prediction (MiRanda score ≥140 and free energy < −18kcal/mol), we only found that only miR-34a-5p putative target sites for the 3’UTR of XBP1 and PD-1 mRNAs (**Table S1**). Therefore, efforts were focused on validating binding of miR-34a to XBP1 (two target sites) and PD-1 (one target site) in NK-cells.

To validate the three putative target sites of miR-34a-5p in the 3’UTR of XBP1 and PD-1 transcripts, we performed a dual luciferase reporter assay (35). We tested the regulatory effect of miR-34a-5p mimics on the expression of *Renilla* Luciferase (Rluc) fused with the predicted target site found in the 3’UTR of XBP1 (**Fig. 2A-B**) or PD-1 (**Fig. 2C**) using HEK293T cells to host the dual luciferase expression vector. For this assay, *Caenorhabditis elegans* miR-67-5p (cel-miR-67-5p), with no putative binding sites found in XBP1 and PD-1 3’UTRs, was used as a negative control to confirm the specific repression of Rluc (carrying candidate target sites) by miR-34a-5p (**Fig. 2A-C**). We then performed a rescue assay by quantifying the expression of Rluc carrying the miR-34a-5p target sites after treatment with miR-34a-5p mimics and rescue with miR-34a-5p antagomir (anti-miR-34a-5p) compared to negative control antagomir (anti-miR NC, a non-targeting negative control for antagomir assays that do not recognize any sequences in human transcriptomes including miR-34a-5p sequence). As expected, normalized Rluc signal was significantly reduced by the presence of miR-34a-5p mimics (plus anti-miR NC; **Fig. S2**) when compared to the rescue condition (plus anti-miR-34a-5p; **Fig. S2**). In addition, we performed a biotinylated miR-34a-5p/mRNA pulldown assay to confirm a significant enrichment of XBP1 and PD-1 mRNAs **(Fig. 2D)**.

**Figure 2:**
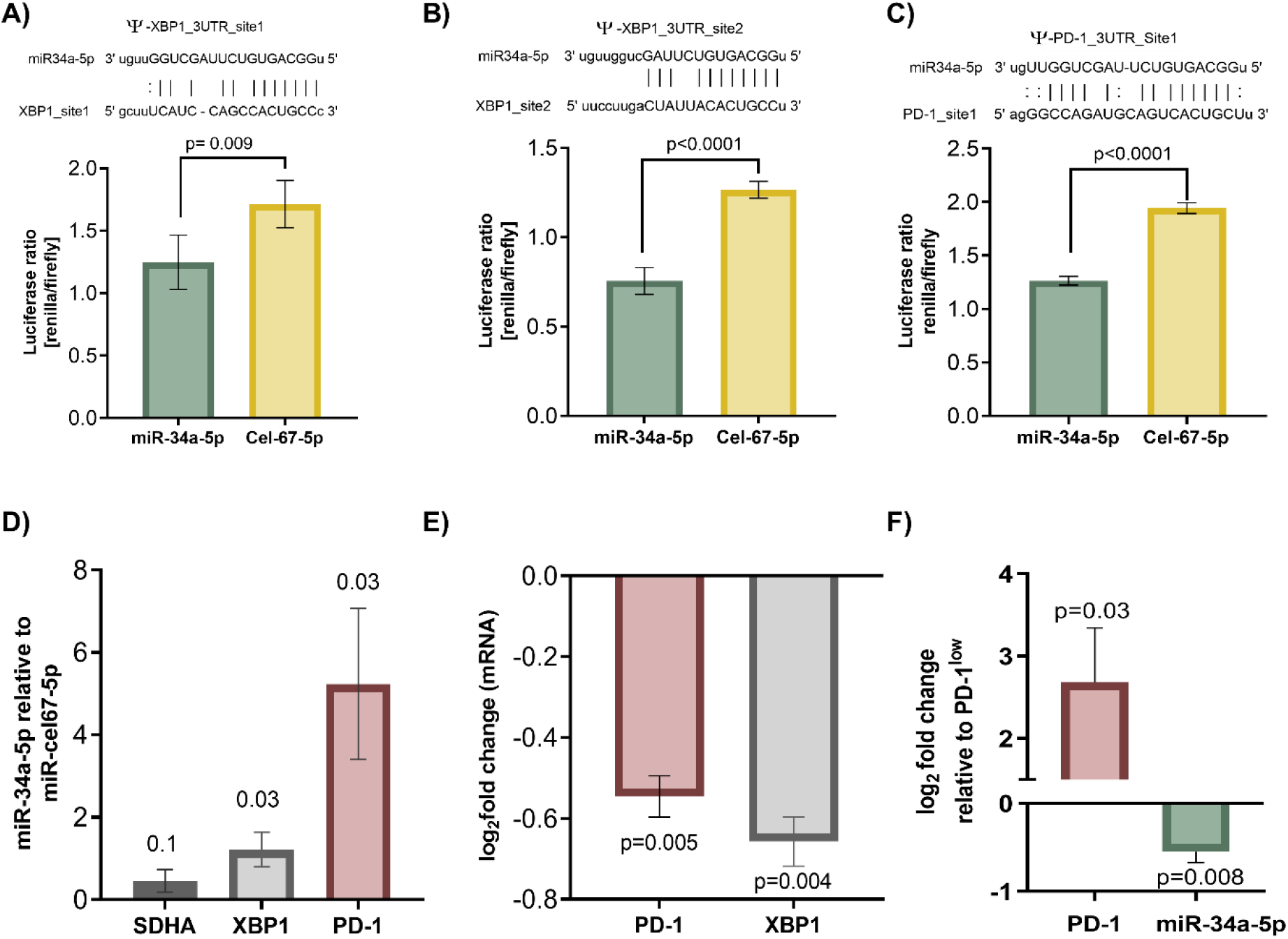
miR-34a-5p directly binds to the 3’UTR of XBP1 and PD-1 transcripts to suppress their expression. miR-34a-5p binding sites, site 1 **(A)** and site 2 **(B)** predicted in the 3’UTR of XBP1 and **(C)** PD-1 were experimentally validated by dual luciferase reporter assay. HEK293T cells co-transfected with miRNA mimics and plasmid construct containing predicted miR-34a-5p binding site showed significant reduction in luciferase signal with miR-34a-5p mimics compared to negative control (cel-miR-67-5p). **(D)** XBP1 and PD-1 mRNAs were significantly enriched in KHYG-1 cell mRNAs pulldown assay using biotinylated miR-34a-5p mimics. *SDHA* gene (non-target) was used as negative control. **(E)** PD-1 and XBP1 mRNA expression in KHYG-1 cell-line was significantly decreased upon miR-34a-5p mimics treatment compared to negative control cel-miR-67-5. A one tailed paired t-test was used for statistical analysis. **(F)** qRT-PCR analysis of miR-34a-5p and PD-1 mRNA in FACS-sorted PD-1^high^ KHYG-1 cells (in contrast to PD-1^low^ KHYG-1 cells) suggests a negative correlation between miR-34a-5p and PD-1 expression (18SrRNA and GAPDH used as endogenous controls for miR-34a-5p and PD-1, respectively). Unpaired t-test two tailed used for statistical analysis.

To further confirm the regulatory effect of miR-34a-5p on XBP1 and PD-1 expression, we treated the PD-1 expressing NK-cell-line KHYG-1 with miR-34a-5p mimics. There was a significant knockdown of XBP1 and PD-1 mRNA in miR-34a-5p mimics-treated KHYG-1 cells compared to negative control treatment (cel-miR-67-5p; **Fig. 2E**). These results confirm that miR-34a-5p can suppress both XBP1 and PD-1 mRNAs in NK-cells. To further confirm the association between miR-34a-5p and PD-1, we assessed the correlative association between miR-34a-5p and PD-1 *in-vitro* by using the PD-1 expressing NK-cell-line KHYG-1. PD-1^high^ and PD-1^low^ KHYG-1 cells were sorted with fluorescent activated cell sorter. Relative quantities of miR-34a-5p and PD-1 transcripts in the two KHYG-1 populations were assessed by qRT-PCR. In agreement with our hypothesis of PD-1 repression by miR-34a-5p, we observed a significant downregulation of miR-34a-5p expression in PD-1^high^ KHYG-1 cells when compared to PD-1^low^ KHYG-1 cells (**Fig. 2F**).

### 2.3 The IRE1α-pathway plays a key role in the downregulation of miR-34a-5p and upregulation of PD-1 expression in NK-cells

A time course experiment for IRE1α blockade using DsiIRE1 (Dicer-substrate RNA interference designed to knock-down IRE1α expression) assay showed significant increase in miR-34a expression in NK-cells that is inversely proportional to IRE1α expression in keeping with an IRE1α negative regulation of miR-34a-5p. For this experiment we again used the PD-1 expressing NK-cell-line KHYG-1 to interrogate the role of IRE1α in the regulation of miR-34a and PD-1. First, we confirmed the inhibitory effect of DsiIRE1 on IRE1α mRNA with maximum knockdown of IRE1α mRNA observed at 12h post-DsiIRE1 treatment (**Fig. 3A**), although IRE1α knockdown lasted for 24h post treatment. As expected, the two arms of miR-34a: miR34a-5p and miR-34a-3p showed an inverted association with IRE1α mRNA expression levels (**Fig. 3B-C**). The expression of both miR-34a-5p and miR-34a-3p peaked at 12h and remained significantly upregulated until 24h post DsiIRE1 treatment. In contrast, the cells with IRE1α inhibited by DsiIRE1 demonstrated significantly reduced PD-1 mRNA expression at 12h post treatment which remained down-regulated until 24h post treatment (**Fig. 3D**). Therefore, we further validated the effect of IRE1α on PD-1 protein expression using TG (IRE1α activator) and DsiIRE1 (IRE1α inhibitor) treatments in the KHYG-1 cells. Our experiments confirmed significant reduction in PD-1 protein expression in the cells treated with DsiIRE1 at 12h and more so at 24h (**Fig. 3E**). On the other hand, IRE1α activation by TG treatment showed a stimulating effect on PD-1 protein expression and this effect was significant even after 24h in the KHYG-1 cells treated with DsiIRE1 compared to negative control DsiNC (**Fig. 3F**).

**Figure 3:**
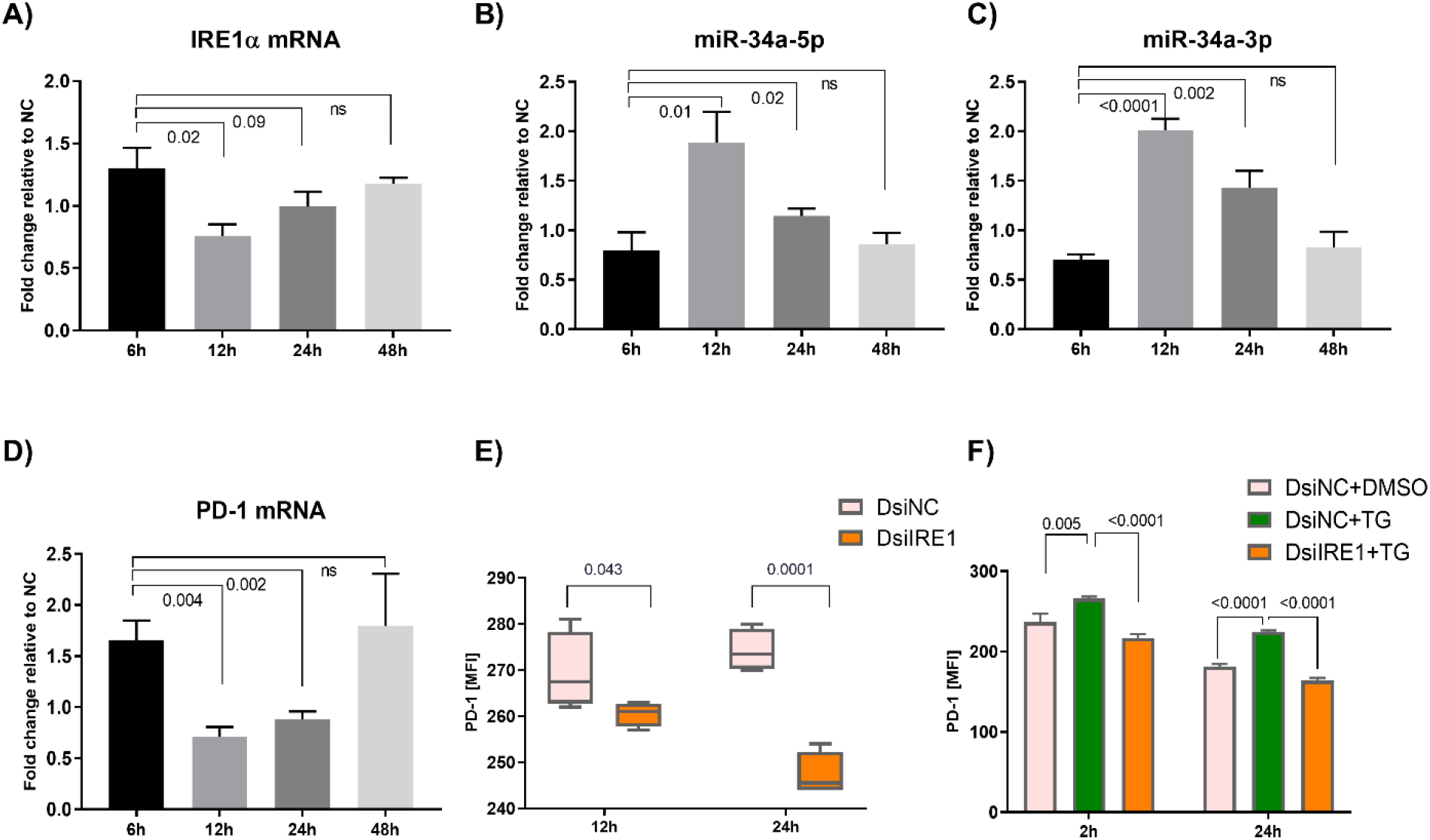
IRE1α suppresses miR-34a expression and increases PD-1 expression in NK-cells. **(A)** Time course experiment for IRE1α blockade by DsiIRE1 in KHYG-1 cells. The highest significant reduction in IRE1α mRNA was observed after 12h of DsiIRE1 treatment compared to negative control DsiRNA (DsiNC) treatment in KHYG-1 cells. The expression of both miR-34a arms: miR34a-5p **(B)** and miR-34a-3p **(C)** was significantly increased in the KHYG-1 cells treated with DsiIRE1. Each miR-34a arm showed an opposite direction to IRE1α mRNA. **(D)** PD-1 mRNA altered in the same direction as IRE1α mRNA. PD-1 mRNA significantly reduced at 12h post DsiIRE1 treatment in KHYG-1 cells and this reduction persisted up to 24h. **(E)** IRE1α blockade significantly reduced PD-1 protein expression in KHYG1 cells after 12h of DsiIRE1 treatment and this effect lasted up to 24h. **(F)** IRE1α activation by TG treatment significantly increased PD-1 expression in KHYG-1 cells when pre-treated with DsiNC but not when IRE1α was downregulated by DsiIRE1. This effect was present in the cells up to 24h after TG treatment.

### 2.4 Activation of the IRE1α-pathway upregulates PD-1 transcription via XBP1s direct binding to target sites in PD-1 promoter regions

We then investigated whether XBP1s putatively binds to the promoter and/or enhancer regions of *PD-1* gene. Putative XBP1 target sites were identified using the canonical XBP1 binding motif as described in JASPAR database (MA0844.1) (36) and a novel variant XBP1 motif that we found in a *de-novo* discovery cohort using XBP1-ChIP assay data previously published (37) (**Fig. 4A**; see methods for details). Next, similar occurrences (at least 70% of similarity) of these two alternative XBP1 motifs were searched in active regulatory regions of *PD-1* previously determined by ATAC-seq (Assay for Transposase-Accessible Chromatin using sequencing) in primary human NK cells (ENCODE; www.encodeproject.org/experiments/ENCSR808HWS/). Two canonical XBP1 target sites were identified in downstream regions of *PD-1* gene while several putative sites were found for the new variant XBP1 motif distributed across the upstream, intronic and downstream regions (**Fig. 4B**).

**Figure 4:**
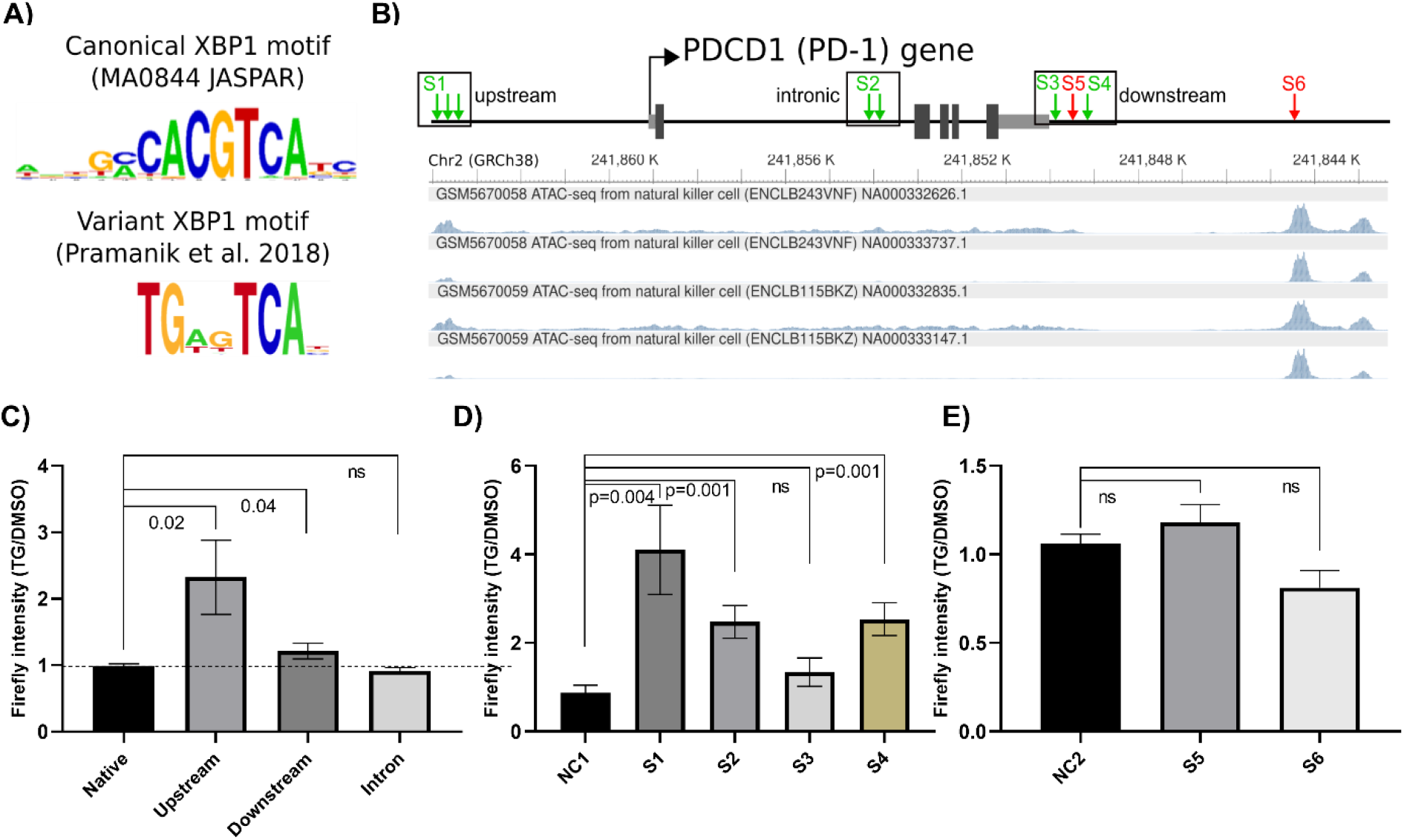
XBP1s regulates PD-1 transcription via target sites in PD-1 promoter regions. **(A)** Two putative XBP1 motifs discovered in regulatory regions bound to XBP1 published elsewhere (37). We identified a XBP1 motif similar to the canonical motif as described in JASPAR database (MA0844) and a novel variant motif not previously published. **(B)** Genomic organization of PD-1 gene showing the location of putative XBP1 target sites. Boxes labelled upstream, intronic and downstream indicate regions selected for experimental validations. Tracks at the bottom of gene diagram depict ATAC-seq data for human NK cells. Red arrows indicate position of canonical XBP1 target sites and green arrow indicate novel variant XBP1 target sites. Dark grey boxes represent exons and light grey boxes the 3’UTRs. **(C)** Multiple XBP1s binding sites were predicted within the upstream, intronic and downstream regions of PD-1. PGL3 vector containing these putative regulatory sequences in the promoter region of a firefly luciferase gene was transfected into HEK293T cells and then treated with TG for activation of XBP1s. PGL3 vectors containing upstream and downstream regulatory regions showed significant increase in firefly luciferase expression upon TG treatment compared to DMSO control. **(D)** Four different binding sites with novel variant XBP1s binding sites (S1, S2, S3 and S4) were cloned into PGL3 vector promoter region. Three binding sites (S1, S2 and S4) showed significant increase in firefly expression upon upregulation of XBP1s (by TG treatment) compared to DMSO treatment as negative control. **(E)** The binding sites with canonical XBP1s motif (S5 and S6) did not elicit upregulation of firefly luciferase upon XBP1s activation (by TG treatment). A one tailed paired t-test was used for statistical analysis.

We selected three putative regulatory regions, upstream, intronic and downstream, of the PD-1 gene where chromatin is accessible in human NK-cells and putative XBP1 binding sites were predicted (**Fig. 4B**). All three regulatory regions were amplified from genomic DNA extracted from KHYG-1 cells and cloned into the promoter of a luciferase gene in a PGL3 vector. HEK293T cells were transfected with PGL3 vectors containing the upstream (810bp), intronic (437bp) or downstream (280bp) regions. Transfected HEK293T cells were then treated with TG (positive control for XBP1s activation) or DMSO (negative control in which XBP1s is not expressed) and the luciferase activity measured using a luciferase reporter assay (compared to native PGL3 vector carrying basic promoter as background control). The results indicate that XBP1s can significantly promote the expression of luciferase gene carrying the upstream and downstream regions containing XBP1s binding sites found in active chromatin regions nearby PD-1 gene (**Fig. 4C)**

We then tested a few selected putative XBP1s binding sites (highest similarity to XBP1s motifs) identified within each regulatory region for further validation. The most abundant XBP1s binding site found in these regions were the variant XBP1s motif (**Fig. 4B**) with the canonical XBP1s motif found only at the downstream region and a more distal accessible region further downstream of PD-1 gene (**Fig. 4B**). Specific putative XBP1s binding sites found in these regulatory regions were cloned into PGL3 vector. The luciferase expression measured in KHYG-1 cells (DMSO or TG treated) transfected with PGL3 vectors carrying candidate XBP1s binding sites revealed that the variant XBP1s binding sites (S1, S2 and S4) can enhance XBP1s-driven expression of luciferase gene (**Fig. 4D**), whilst the canonical XBP1 binding sites (S5 and S6) were not able to elicit any significant change in luciferase expression (**Fig. 4E**).

### 2.5 The IRE1α-pathway is impaired and associated with reduced effector function in primary NK cells of patients with Hodgkin lymphoma

To investigate the translational relevance of our findings, our study was extended to primary NK-cells (pNK-cells) obtained from patients with Hodgkin Lymphoma (HL), a blood cancer in which NK-cells expressing PD-1 are known to be functionally impaired (26). We tested *ex-vivo* non-expanded pNK-cells taken from peripheral blood from patients with HL prior to therapy, to establish the base levels of *ex-vivo* IRE1α-endonuclease activity. XBP1s expression was quantified by flow cytometry in HRS-stimulated pNK-cells from patients with HL and compared to age-matched healthy participants. Mononuclear CD3^-^CD14^-^CD19^-^CD56^+^ pNK-cells were enumerated and classified into conventional pNK-cell subsets based on CD56 and CD16 expression: CD56^bright^CD16^-ve^ and CD56^dim^CD16^+^. Following HRS-stimulation, XBP1s expression was reduced in both CD56^bright^CD16^-ve^ (**Fig. 5A**) and CD56^dim^CD16^+^ subsets (**Fig. 5B**) in HL patients, indicating that IRE1α-endonuclease activity (required to effectively process the functional XBP1s isoform) is reduced in pNK-cells from cancer patients.

**Figure 5:**
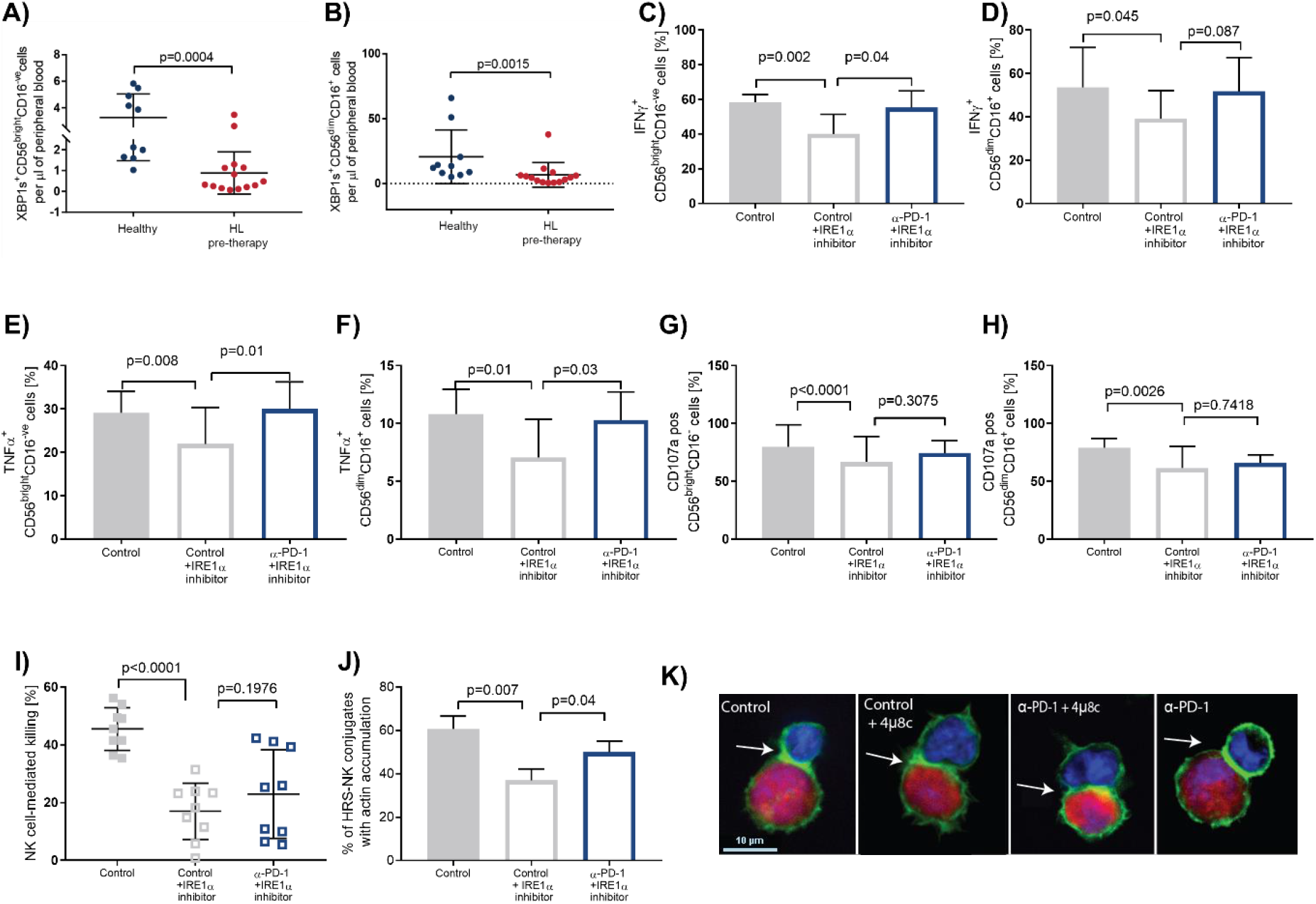
IRE1α-pathway is impaired in primary NK-cells from patients with Hodgkin Lymphoma and PD-1 blockade partially restores the IRE1α-pathway and pNK-cell effector functions. The effect of Hodgkin Reed-Sternberg (HRS) cells on IRE1α-pathway activation was tested in pre-therapy primary NK-cells (pNK-cells) from patients with Hodgkin Lymphoma (HL) compared to age-matched healthy participants. PBMCs were incubated for 4 hours in the presence of HRS-cells (KMH2) admixed at 1:1 ratio. **(A)** XBP1s expression in the CD56^bright^CD16^-ve^ NK-cell subset was quantified by flow cytometry data and found to be downregulated in pre-therapy pNK-cells from patients with HL (compared to pNK-cells from healthy participants). **(B)** The absolute numbers of XBP1s^+^ CD56^dim^CD16^+^ NK-cell subset was quantified by flow cytometry showing downregulation in pre-therapy pNK-cell from patients. Mann-Whitney test (n=14) used for statistical analysis. **(C-I)** The effector functions of NK-cells from patients with HL were reduced by IRE1α inhibitor (4μ8c 60μM) when compared to Control (DMSO and IgG_4_ isotype), after stimulation with HDLM2 cells for 6 hours. PD-1 blockade (α-PD-1) partially restored pNK-cell functions. Cytokines **(C, D)** IFNγ and **(E, F)** TNFα were assessed by FACS in CD56^bright^CD16^-ve^ and CD56^dim^CD16^+^ cells and were significantly reduced after IRE1α inhibition. PD-1 blockade reversed IFNγ release reduced by IRE1α-inhibition in CD56^bright^CD16^-ve^ subset of NK-cells. TNFα release reduced by IRE1α-inhibition was reversed by PD-1 blockade in both CD56^bright^CD16^-ve^ or CD56^dim^CD16^+^ NK-cells subsets. A Paired t-test was used for statistical analysis. At the end of a two-week expansion NK-cells from patients with HL were assessed. **(G, H)** Cells were stained with CD107a antibody to assess the level of de-granulation in NK-cell subsets. IRE1α-blockade inhibited de-granulation however PD-1 blockade did not reverse IRE1α blockade-mediated impairment of NK-cell de-granulation in (**G**) CD56^bright^CD16^-ve^ or (**H**) CD5^dim^CD16^+^subset of NK-cells. (**I**) IRE1α-blockade inhibited lysis of HRS target cells but no significant difference in cytotoxicity was observed upon PD-1 blockade. Statistical differences between experimental groups were evaluated by Student’s t-test (n=3). **(J)** NK-cell immune synapse formation between NK-cell and HRS cells that is impaired by IRE1α-blockade is restored after PD-1 blockade with increased F-actin polymerization at the NKIS. Statistical differences between experimental groups were evaluated by unpaired t-tests. Histograms show the mean ± SD from 3 independent experiments with 30 conjugates analysed per experiment. **(K)** Representative images of NK-cell immune synapse formation between NK-cell (DAPI, blue) and HRS cells (APC, red) showing the expanded NK-cells from patients with HL stimulated with HDLM2 for 6 hours in the presence of IRE1α inhibitor, PD-1 blockade (α-PD-1) + 4μ8c. Arrows indicate actin accumulation (green) at the NK-cell/HRS synapse site. In the control assays, HRS target cells and pNK-cells were co-incubated with immunoglobulin (Ig) G4 isotype control. Original magnification ×63.

Next, we evaluate if there is still a remnant role for the impaired IRE1α-pathway in the regulation of pNK-cell effector functions in HL. As the numbers of intratumoral pNK-cells in patients with HL are limited, circulating pNK-cells from pre-therapy blood samples in patients with HL were expanded *in-vitro*. Then the functional impact of IRE1α-blockade (using IRE1α-inhibitor 4μ8c) was tested in the presence of HDLM2 target cells. This showed a reduction of IFNγ (**Fig. 5C,D**) and TNFα cytokines release (**Fig. 5E,F**), CD107a de-granulation (**Fig. 5G,H**), HRS target cell killing (**Fig. 5I**) and F-actin accumulation at the NK-cell immune synapse (NKIS; **Fig. 5J,K**) in pNK-cells from patients with HL.

These results strongly suggest that IRE1α-pathway still retains a residual activity and thus it may be possible to restore a ‘healthy’ homeostasis. Considering our previous study demonstrates pNK-cell effector function are significantly improved by the immunotherapy medication Pembrolizumab (monoclonal antibody designed to target and block PD-1) (26), we sought to determine whether PD-1 blockade restores NK-cell effector functions that are reduced due to impairment of IRE1α-pathway. Cytokine secretion (IFNγ, TNFα), de-granulation (CD107a), HRS target cell killing and F-actin accumulation at the NKIS were tested in pNK-cells from patients with HL. We tested the effect of Pembrolizumab (α-PD-1) on pNK-cell functions upon IRE1α-inhibition (**Fig. 5C-K**). PD-1 blockade restored both IFNγ (**Fig. 5C,D**) and TNFα (**Fig. 5E,F**) cytokines release in pNK-cells (except for IFNγ in CD56^dim^CD16^+^). However, PD-1 blockade did not restore de-granulation (**Fig. 5G,H**) and cytotoxicity (**Fig. 5I**) functions, despite of a significant increase in immune synapse formation when PD-1 is blocked (**Fig. 5J-K**), in pNK-cells from patients with HL. Moreover, we observed that pNK-cells from patients treated with Pembrolizumab alone have restored IFNγ release (**Fig. S3A,B**) and immune synapse (**Fig. S3H,I**) but none of the other effector functions were enhanced (**Fig. S3**) suggesting the improvement in NK-cell function induced by PD-1 blockade is only partially by rescuing IRE1α-pathway impairment.

In confirmatory experiments, we also measured cytokine secretion, de-granulation and cytotoxicity upon IRE1α-blockade on the SNK10 cell-line stimulated with K562 and HDLM2 target cells. The results showed significant reduction of all effector functions including IFNγ release (**Fig. S4A,B**), TNFα release (**Fig. S4C,D**), CD107a de-granulation (**Fig. S4E**) and lysis of target cells (**Fig. S4F**) confirming the IRE1α-pathway has an important role in NK-cell effector function. Moreover, IRE1α-blockade efficiently inhibits XBP1 splicing in NK-cells when exposed to all three tested target cells (K562, KMH2 and HDLM2; **Fig. S4G**), further confirming the IRE1α-pathway involvement in XBP1 splicing in NK-cells. We also tested if the IRE1α-pathway has a role in NK-cell movement and synapse formation. SNK10 cells motility was assessed by live cell imaging (**Fig. S4H)**. IRE1α-blocked NK-cells travelled shorter distances and at lower velocity (**Fig. S4H)**.

Since a recent study has reported that NK-cells can acquire functional PD-1 transferred from cancer cells via trogocytosis (34), we also investigated whether the PD-1 expression detected in the cellular membrane of NK-cells used in our assays could have been acquired from malignant HRS cells due to trogocytosis in HL instead of an endogenous expression in NK-cells. First, PD-1 expression was tested in a range of HL cell-lines and none of the tested lines expressed PD-1 mRNA or protein (**Fig, S5A-C**). Then, we tested the expression of PD-1 and CD30 (a marker of HRS cells) on pNK-cells in pre-therapy blood samples in patients with HL. The results demonstrate that pNK-cells express PD-1 but not CD30 (**Fig. S5D**) supporting the notion that PD-1 expression on pNK-cells is not due to trogocytosis but an endogenous expression in pNK-cells.

### 2.6 miR-34a-5p is negatively correlated to PD-1 in NK-cells from patients with Hodgkin lymphoma

Having established that the IRE1α-pathway is impaired in the pNK-cells of patients with HL, we next compared the native expression of miR-34a-5p and PD-1 in pNK-cells from 9 healthy individuals and 11 patients with HL. Our results with NK-cell lines indicate that miR-34a-5p negatively regulates PD-1 expression, therefore an inverse relationship is expected to emerge between miR-34a-5p and PD-1 expression in pNK-cells. We used a PrimeFlow RNA assay to simultaneously measure expression of miR-34a-5p and PD-1 protein by flow cytometry. With the specific oligonucleotide target probe, the assay hybridises and amplifies miRNA molecules at single-cell resolution without the requirement for RNA extraction and quantification by conventional qRT-PCR. We observed two distinct clusters based on miR-34a-5p and PD-1 expression in pNK-cells from patients with HL (Cluster 1: CD56^+^CD3^-ve^PD-1^+^miR-34a-5p^low^ and Cluster 2: CD56^+^CD3^-ve^PD-1^+^miR-34a-5p^high^; **Fig. 6A**). We then tested the two sub-groups for correlation between miR-34a-5p and PD-1 expression with CD56^+^CD3^-ve^PD-1^+^miR-34a-5p^high^ sub-group showing significant negative correlation (Pearson’s correlation coefficient R = −0.92; p = 0.009; **Fig. 6A**) whilst the CD56^+^CD3^-ve^PD-1^+^miR-34a-5p^low^ sub-group showed negative correlation, however this did not reach statistical significance (Pearson’s correlation coefficient R = −0.83; p = 0.08; **Fig. 6A**). No significant negative correlation was found in healthy subjects with detectable levels of PD-1 expression (Pearson’s correlation coefficient R = 0.18; p = 0.73; data not shown). This inverted relationship confirms our observations in NK-cell lines in which miR-34a-5p is identified as a key inhibitory regulator of PD-1 expression.

**Figure 6:**
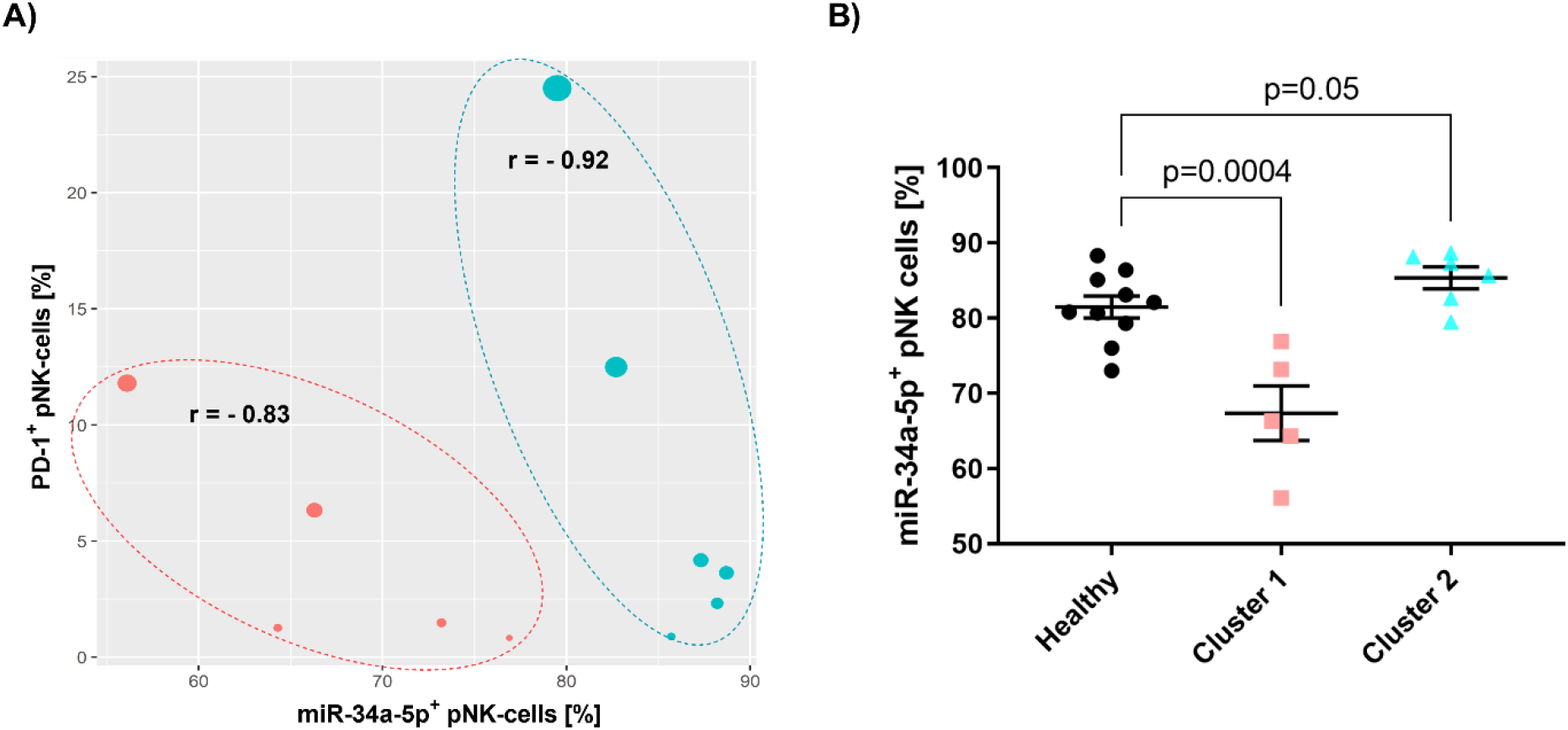
Inverse correlation between miR-34a-5p and PD-1 expression in primary NK-cells from patients with Hodgkin lymphoma. **(A)** Simultaneous quantification of miR-34a-5p and PD-1 protein in *ex-vivo* pNK-cells from HL patients showed significant negative correlation between miR-34a-5p and PD-1 protein expression levels in two sub-groups CD56^+^CD3^-ve^PD-1^+^miR-34a-5p^low^ (Cluster 1) and CD56^+^CD3^-ve^PD-1^+^miR-34a-5p^high^ (Cluster 2). Size of the dots indicates combined miR-34a-5p and PD-1 protein expression levels. CD56^+^CD3^-ve^PD-1^+^miR-34a-5p^high^ sub-group showed a significant negative correlation (p = 0.009) whereas the negative correlation in the CD56^+^CD3^-ve^PD-1^+^miR-34a-5p^low^ sub-group did not reach significance (p = 0.08). K-means clustering was used to group samples based on miR-34a-5p and PD-1 expression. Pearson’s correlation test was used for statistical analysis. **(B)** miR-34a-5p expression in patients with HL from Cluster 1 is significantly down regulated compared to healthy individuals whereas patients with HL from Cluster 2 have a marginally significant increase in miR-34a-5p expression level.

Intriguingly, we also found miR-34a-5p expression is reduced in pNK-cell Cluster 1 (CD56^+^CD3^-ve^PD-1^+^miR-34a-5p^low^ sub-group) compared to healthy individuals but not in Cluster 2 (CD56^+^CD3^-ve^PD-1^+^miR-34a-5p^high^) (**Fig. 6B).** Arguably, the impaired IRE1α-pathway observed in pNK-cells from patients with HL may also impact the regulation of miR-34a-5p expression and consequently PD-1 expression. However, a more detailed understanding of the mechanistic nature of IRE1α-pathway impairment in cancer will require further studies with an appropriate design to address these specific clinical questions.

## 3 Discussion

Herein we uncovered a new mechanistic association between the miRNA miR-34a-5p, transcription factor XBP1 and immune checkpoint PD-1 upon IRE1α activation to modulate human NK-cell effector function and tolerance. In the presence of cancer cells, NK-cells activate the IRE1α-pathway, triggering a rapid change in XBP1s-mediated gene regulation. This is amplified by a reduction of miR-34a-5p which in turn represses XBP1 (as well as PD-1) expression. Moreover, XBP1s promotes PD-1 expression that is further enhanced by the IRE1α-mediated reduction of miR-34a-5p, thus fine-tuning NK-cell effector function. IRE1α activation is impaired in NK-cells within patients with HL, resulting in a disruption of XBP1/miR-34a axis and PD-1 expression within NK-cells of patients with HL.

Several studies indicate that the NK-cell/tumor cell PD-1/PD-L1 axis is involved in immune evasion in blood cancers including HL (25–27), however the molecular mechanisms that fine-tune PD-1 expression are unknown. The findings from the current study indicate that the IRE1α-pathway regulates PD-1. Upon initial contact with cancer cells (i.e., malignant HRS cells), the IRE1α-pathway is induced resulting in upregulation of NK-cell effector function. However, under sustained activation PD-1 expression is upregulated, promoting immune tolerance (or exhaustion). Arguably, this ‘negative feedback’ serves as a regulatory mechanism to adjust to chronic NK-cell activation imposed by cancer-induced stress. The second outcome of IRE1α-pathway activation is to fine-tune PD-1 expression by removing the inhibitory miR-34a-5p to increase PD-1 expression (**Fig. 7**). Chronic activation of the IRE1α-pathway downregulates miR-34a-5p, which increases PD-1 expression and its inhibitory effects in NK-cells. In line with these findings, miR-34a-5p and PD-1 were inversely correlated in pNK-cells of patients with HL, implicating the IRE1α/miR-34a-5p/PD-1 axis as a novel mechanism in NK-cell linking immune cell activation and tolerance through IRE1α and immune checkpoint PD-1.

**Figure 7:**
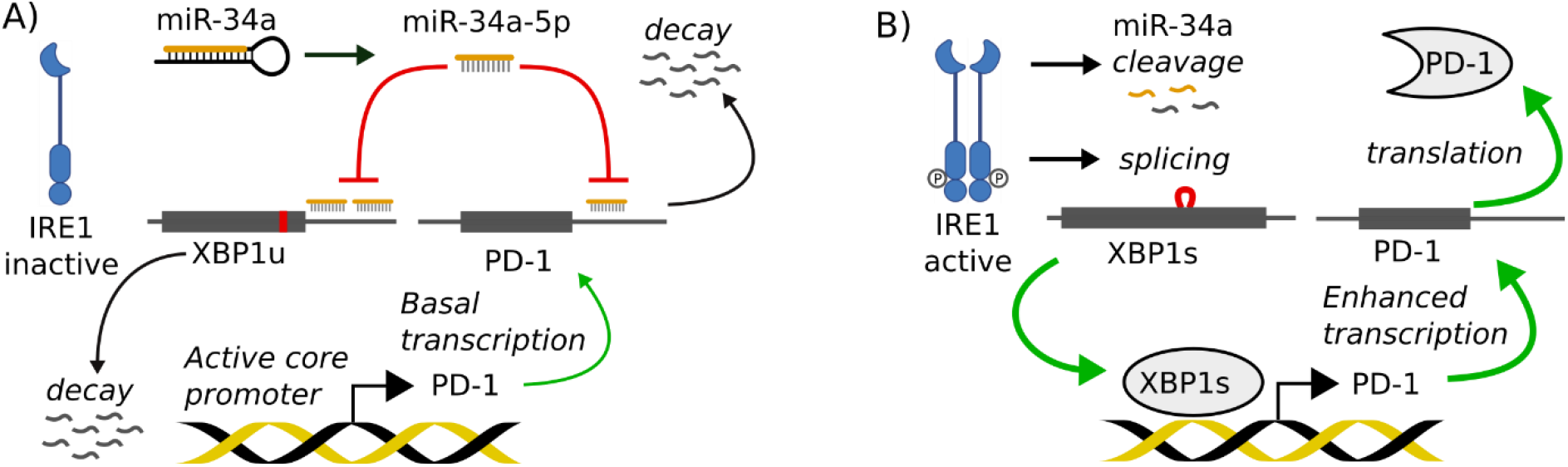
Proposed mechanism for the dual role of IRE1α-pathway in regulating the XBP1/miR-34a axis to increase PD-1 expression. **(A)** When IRE1α is inactive, the XBP1 transcript is not spliced (XBP1u), which lacks a transcription factor domain and is rapidly degraded by proteasome activity. In IRE1α inactive state, the miR-34a biogenesis is not disrupted by IRE1α endonuclease activity and functional miR-34a-5p is produced to suppress XBP1u expression further increasing XBP1u decay by RNA interference. The expression of miR-34a-5p also plays a role to maintain PD-1 expression under minimal thresholds. **(B)** Upon IRE1α activation, miR-34a-5p is degraded by IRE1α-endonuclease by cleavage of the primary miR-34a hairpin (via CUGCAG motif); through a similar mechanism a 26nt intron is removed from XBP1 mRNA encoding an effective transcription factor (XBP1s) that changes the transcription of several downstream genes involved in immune response such as PD-1 which is now released from miRNA suppression and can be effectively translated into functional PD-1 protein.

Recent studies have demonstrated IRE1α as a “double-edged sword” that co-opts its endonuclease activity to regulate the activation of XBP1s while simultaneously repressing a subset of miRNAs [11]. IRE1α endouclease activation causes rapid decay of CUGCAG motif-containing miRNAs (i.e., miR-17, miR-34a, miR-96 and miR-125b), and hence strongly increases the expression of several genes affecting multiple cellular pathways [11, 42]. The current study is the first to provide experimental evidence that IRE1α works as a dual regulatory switch to enhance PD-1 expression in NK-cells through simultaneous activation of XBP1s and repression of miR-34a-5p.

Interestingly, miR-34a-5p and PD-1 expression are negatively correlated in pNK-cells from patients with HL, suggesting an *in-vivo* regulatory role of miR-34a-5p in the PD-1/PD-L1 immune checkpoint axis. Neither PD-1 mRNA or protein was expressed across HL cell-lines, nor was the HRS marker CD30 observed in pNK-cells in patients with HL, indicating that the expression of PD-1 in the NK-cells of patients cannot be explained by trogocytosis as previous published [34]. The IRE1α/miR-34a/PD-1 axis may be one factor to explain the well-documented contradictory findings regarding PD-1 expression on NK-cells [25, 32, 44, 45]. Notably, miR-34a-5p has been previously found to be a master regulator of tumor suppression [46], and to enhance the repression of PD-L1 in acute myeloid leukemia and lung cancer [13, 14]. Therefore, the IRE1α/miR-34a regulatory circuitry in the PD-1/PD-L1 immune checkpoint axis may partially explain the immune exhaustion commonly observed in blood cancers [43]. We also found that PD-1 blockade only partially reversed NK-cell dysfunction and neither de-granulation nor cytotoxic function was restored by PD-1 blockade after IRE1α-pathway inhibition. Hence, our findings support the current growing interest in small RNA-based therapies to manipulate immune checkpoints in immune cells including NK-cells to improve cancer immune therapies in blood cancers [47, 48].

In conclusion, a hitherto undescribed NK-cell dual regulatory axis is outlined, by which the IRE1α-pathway fine-tunes NK-cell effector activity through an inter-connected activation of the transcription factor XBP1s and inhibition of miR-34a-5p to modulate PD-1 expression. IRE1α activation is impaired in NK-cells within patients with HL. Hence the dual role IRE1α-pathway plays in regulating the XBP1/miR-34a axis and PD-1 expression within NK-cells is disrupted in patients with HL. Put together, the findings may help explain previous conflicting findings regarding PD-1 expression in NK-cells and assist the development of new immunotherapeutic strategies.

## 4 Materials and Methods

### 4.1 NK-cell stimulation using cell-lines

NK-cell-lines used were SNK10 (chronic active Epstein-Barr virus) and KHYG-1 (NK-cell leukemia). Target cell-lines were HDLM2 (HL), KMH2 (HL), K562 (HLA-deficient erythroleukemia). Effector:Target (E:T) ratios used in experimental assays were 1:1. NK-cell lines were typically stimulated for 2 hours (K562) or 8 hours (KMH2 and HDLM2).

### 4.2 IRE1α-pathway activation and inhibition assays

Activation of the IRE1α-pathway using thapsigargin (TG) and inhibition of the IRE1α-pathway using small molecule inhibitors 8-formyl-7-hydroxy-4-methylcoumarin (4μ8c) or 6-Bromo-2-hydroxy-3-methoxybenzaldehyde (6-bromo). Knock-down of IRE1α expression were performed using Dicer-substrate short interfering RNA (DsiRNA; Integrated DNA technologies) as outlined in the supplemental material.

### 4.3 Identification and validation of miRNA target gene regulation

Putative target sites in the 3’UTR region of XBP1 and PD-1 were predicted for IRE1α-regulated miRNAs (miR-17, miR-34a, miR-96 and miR-125b) using miRanda algorithm (38). Mature miRNA sequences for all four miRNAs were downloaded from miRBase database (Release 22) (39) and the XBP1 and PD-1 transcript sequences were retrieved from GENCODE Release 21 (Reference Human Genome GRCh38) (40). Only predicted target sites with stringent parameters (free energy ≤ −18 and miRanda score ≥ 140) were considered for further experimental validations (**Table S1**). Functional validation of miRNA binding sites including the dual luciferase assay and transfection of miRNA mimics is outlined in more details in the supplemental material.

### 4.4 Identification and validation of XBP1 target sites in PD-1 gene

The computational prediction of XBP1 binding sites was performed on regulatory regions of the PDCD1 (PD-1) gene found to be active in human Natural Killer cells (ENCODE; www.encodeproject.org/experiments/ENCSR808HWS/). We used MEME motif discovery program (41) for a *de novo* discovery of XBP1 binding sites on sequences enriched in regions experimentally determined by XBP1-ChIP assay and RNA-seq (as described elsewhere (37) and datasets available at ArrayExpress: E-MTAB-6327, E-MTAB-6894 and E-MATB-7104). We identified over-represented DNA patterns (or motifs) in the XBP1 binding regions of the top 500 most upregulated genes in activated TH2 (high levels of XBP1s) versus naïve TH2 (889 peaks) and the top 500 most down-regulated genes in activated TH2 versus naïve TH2 (595 peaks). Two putative XBP1 motifs were found: one canonical motif similar to the one described in JASPAR database (MA0844) and a novel variant XBP1 motif not yet described in any available database or publication. TAMO package (42) was used for the identification of putative XBP1 binding sites in the promoter regions of PD-1 gene. Only putative binding sites at least 70% similarity to canonical or variant XBP1 motifs were selected.

A luciferase reporter system (pGL3 basic vector, Promega) was used to validate the direct interaction of XBP1s and the predicted promoter regions of PD-1 as well as the individual XBP1s binding sites identified within PD-1 promoter regions. More details of methods are described in supplemental material.

### 4.5 Patient samples

Pre-therapy blood samples from 14 patients with HL were collected before therapy (median age 37 years, range 16-73; F:M ratio 9:5; 6 stage I-II, 6 stage III-IV, 2 stage unknown; 10 nodular sclerosing, 1 lymphocyte rich, 3 classical HL otherwise not classified); and from 10 healthy participants (median age 37 years, range 25-48, F:M ratio 5:5). Peripheral blood mononuclear cells (PBMC) were isolated and cryopreserved, as described previously (43). This study conformed to the Declaration of Helsinki, and informed consent was provided by all participants in accordance with participating hospitals/research institute Human Research Ethics Committee guidelines.

### 4.6 NK-cell functional assays

pNK-cells were isolated from cryopreserved PBMCs and expanded as previously described (26). Flow cytometry, NK-cell migration, NK-cell immune synapse (NKIS) formation and cytotoxicity are outlined in the supplemental. For PD-1 blockade experiments, NK-cells were cultured for 72 hours with 10μg/ml of pembrolizumab™ (or IgG_4_ control), prior to the addition of targets. At the completion of the culture the KHYG-1 cells or pNK-cells were collected and incubated with an equal number of targets. Assays were performed in triplicate. More details provided in supplemental material.

### 4.7 Statistical analysis

All statistical analysis was done using GraphPad Prism 7.00 (GraphPad Software). Comparison of measurements was performed using paired 2-tailed T tests (except when stated otherwise in figure legends). Only P < 0.05 was considered significant.

## Author contributions

Conception and design: A.S. Cristino, M.K. Gandhi, S. Mujaj, J. Nourse, K. Bednarska, G. Thillaiyampalam

Financial support: A.S. Cristino, M.K. Gandhi

Administrative support: M.K. Gandhi

Provision of patient samples: M.K. Gandhi

Collection and assembly of data: A.S. Cristino, K. Bednarska, G. Thillaiyampalam, M. B. Sabdia, S. Mujaj, J. Nourse, L. Meridia de Long, J. Collins, F. Vari, Q. Cui, J. Gunawardana

Data analysis and interpretation: A.S. Cristino, K. Bednarska, G. Thillaiyampalam, M.K. Gandhi, S. Mujaj, J. Nourse

Manuscript writing: All authors

Final approval of manuscript: All authors

Accountable for all aspects of the work: All authors

## Funding

Supported by the Leukaemia Foundation and the Mater Foundation (M.K.G); Centre for Children’s Immunotherapy Research Grant, ID 50324 (M.K.G and A.S.C); Princess Alexandra Hospital Foundation (M.K.G. and A.S.C); University of Queensland PhD scholarships for international students (G.T. and K.B), and a Queensland Health Junior Doctor Research Fellowship (J.C). The Translational Research Institute is supported by the Australian Government.

## Notes

**Conflict-of-interest**, M.K.G. has received research funding from Beigene and Janssen. There are no other relevant competing financial interests.

### Competing Interest Statement

The authors have declared no competing interest.

