## Supplemental material for "The IRE1α-endonuclease regulates PD-1 expression through a novel XBP1/miRNA-34a axis within Natural Killer cells"

**This PDF file includes:**

Methods Page 2

Figure S1 Page 10

Figure S2 Page 11

Figure S3 Page 12

Figure S4 Page 14

Figure S5 Page 16

Table S1 Page 18

References Page 18

### Methods

#### Flow cytometry assays

All antibodies used were human specific. These were: XBP1s, CD3, CD14, CD19, CD56 and CD16 (BD Biosciences), and CD107a, IFNγ and TNFα (BioLegend). Following staining, flow cytometry was performed using the LSR II™ (BD Biosciences) and analyzed using FlowJo software (Tree Star Inc.). To exclude dead cells each panel included a live/dead cell detection step (Zombie NIR Fixable Viability kit, BioLegend). To detect CD107a surface expression, SNK10 cells were incubated with cell trace violet-stained target cells at 1:1 ratio for 1 hour. CD107a-APC antibody (1/200 dilution) and monensin (1/1500 dilution) were then added to each well and cells were incubated for further 5 hours. IFNγ, CD107a degranulation, NK-cell assays were performed as previously described (1, 2).

To assess the PD-1 protein expression on KHYG-1, PD-1 PE (clone PD1.3.1.3, Miltenyi Biotec) was added to KHYG-1 and stained for 20 minutes on ice. The cells were then acquired using BD Fortessa x20 flow cytometer controlled by FACSDiva software (BD Biosciences). Data analysis was performed in FlowJo 10 software (Tree Star). Acquired cells were initially defined by forward and side-scatter and then gated for PD-1 expression relative to an unstained control.

The Prime flow RNA assay kit (Thermo Fisher Scientific) was used to detect miR-34a-5p in pNK-cells expanded from patients with HL. The assay is an in-situ hybridisation flow cytometry assay with single-cell resolution, enabling simultaneous detection of RNA targets and proteins using fluorochrome-conjugated antibodies. The detection of microRNA was performed using PrimeFlow™ microRNA Pre-treatment Buffer as per manufacturer's protocol.

#### NK cell migration assay

Live cell imaging of untreated or IRE1 inhibited (60µM of 4µ8c) SNK10 cells was used to assess changes in motility (3). For this, cell trace far red stained NK-cell-lines were seeded onto 6-well slides coated with HeLa cells and images were taken every two minutes for five hours using the Olympus xCellence real-time live cell imaging system. Tracking of cells was done using the mTrackJ plugin for image J. For this, 20 cells were randomly chosen and tracked. The mean distance and the mean velocity of cells were used for analysis.

#### NK-cell cytotoxicity assay

To analyze NK-cell cytotoxicity of target cells, K562 or HDLM2 target cells were stained with Carboxyfluorescein succinimidyl ester (CFSE) dye (Sigma-Aldrich). SNK10 cells were incubated with CFSE stained target cells at a 1:1 effector: target ratio. For each experiment target controls, with no effector cells (pNK or SNK10 cells), and effector controls, with no target cells (K562 or HDLM2 targets), were included.

NK effector cells were incubated with their targets at 37°C with 5% $CO_2$ for 3 hours unless stated otherwise. After this time cells were stained for expression of markers of interest as described below. After staining 10,000 CountBright counting beads (Invitrogen) were added to each well in 100µL of PBS supplemented with 2% FCS to calibrate numbers. Flow cytometry was performed using the FACSCanto™ (BD Biosciences) and a minimum of 5,000 events were collected in the beads gate.

The number of cells in each gate was determined using FlowJo software (Tree Star Inc.). Briefly, beads (B) and cells were gated based on forward scatter (FSC) and side scatter (SSC). The cells were then further divided into CFSE positive target cells (T) and CFSE negative effector cells (E). To assess the percent of target cell lysis the following formula was used: Lysis = 1- ($T_{exp}$ / B

÷ Average ($T_{con}$ / B)) where $T_{exp}$ represents the number of target cells in experimental wells and $T_{con}$ represents the number of target cells in control wells.

#### NK-cell immune synapse analysis

KHYG-1 or FACS-sorted $CD3^{-}CD56^{+}$ pNK-cells (1.5x$10^{5}$) were mixed with HDLM2 cells (1.5x$10^{5}$) (pre-stained with cell tracker red, Thermofisher Scientific) and incubated in FACS tubes at 37 °C and 5% $CO_2$ for 15 min. Conjugates were then transferred to silane-coated microscope slides (Electron Microscopy Sciences) and incubated for another 15 min at 37 °C. After a final incubation of 30 min., cells were washed with PBS (Life Technologies) and fixed and permeabilized using 4% para-formaldehyde for 20 min at room temperature. After fixation, coverslips were permeabilized in 0.25% Triton-X-100 for 5 min. Blocking and antibody dilution were performed in PBS/0.2% NaAzide/0.25% BSA. F-actin was stained with Alexa Fluor 488 phalloidin (Life Technologies) for 30 min. At final stage cells were stained with DAPI for 10 min. Cells were mounted using Menzel-Glaser 22 mm x 40 mm coverslips (Fisher Scientific) and buffered glycerol with anti-fade mountant (Harvard OMX). Between the staining repetitive washing steps were performed with PBS.

Confocal microscopy was performed using the Nikon spinning disc confocal microscope. Images were collected using the Nikon Plan Fluor 40X objective (NA = 1.3). The NIS-Elements software was used for image acquisition. Image sets to be compared were acquired during the same session and using the same acquisition settings.

Quantitative image analyses of conjugates were performed using ImageJ analysis software. Quantification of F-actin polarization at the immune synapse was based by random selection of 30 conjugate images containing an APC–stained HDLM2 cell in contact with a NK-cell. To quantitate recruitment of F-actin to the immunological synapse, polarization of actin at the NK-cell contact site was measured and then scored based on the median thickness of actin for untreated cells. Those conjugates showing a distinct actin band at the immune synapse site were considered polarized (above median for untreated cells, score=1). Conjugates with weak polymerization (actin thickness below median for untreated cells, score=0) or showing moderate polymerization (score=0.5). Data were plotted using the Prism 7.0 (GraphPad) software. Statistical significance was calculated using a Student's *t*-test (two-tailed).

#### RNA extraction and cDNA synthesis

For RNA extractions, following stimulation with target cells, NK-cells were separated from remaining targets using the NK-cell isolation kit (Miltenyi Biotec), as per the manufacturer's instructions. These stimulated NK-cells were >90% pure by flow cytometry. RNA was isolated from cells using the RNeasy Mini kit (Qiagen) and following the manufacturer's protocol. RNA was reverse transcribed to cDNA using SuperScript® III Reverse Transcriptase (Life Technologies). Freshly synthesized cDNA was diluted to 2.5ng/µL of the starting RNA concentration using nuclease-free water.

#### Quantitative RT-PCR of XBP1s and miRNA expression

XBP1 splicing was determined using quantitative PCR (qPCR) and primers that spanned the splice site. qRT-PCR was performed with the PowerUp SYBR Green Master Mix (Applied Biosystems by Life Technologies) using 5ng of cDNA input. Quantification of samples (relative to controls) was determined using QuantStudio Real-Time PCR software (Life Technologies) relative quantification analysis. For all qPCR analysis, untreated SNK10 cells were used as the control unless stated otherwise. Relative expression values given were normalized to the housekeeping gene; *GAPDH,* to account for slight variations in cDNA concentrations. Experiments were repeated three times or more with technical triplicates and no-treatment negative controls.

Total RNA was extracted from NK-cell-lines SNK10 or KHYG-1 and HEK293T cells using miRNeasy Kit (Qiagen) and cDNA synthesised and amplified (input cDNA 2.5ng/µL) using Qiagen miScript PCR system which includes miScript II RT Kit and SYBR Green PCR Kit. miRNA expression was quantified on LightCycler® 480 Real-Time PCR System (Roche Molecular Systems) or ViiA™ 7 Real-Time PCR System (Thermo Fisher Scientific). Comparative quantification between miR-34a-5p and housekeeping genes RNU6 or 18srRNA was used to determine the relative expression delta Ct and then calibrated to an appropriate control sample.

#### IRE1α inhibition and knock-down assay

Inhibition of IRE1α-mediated splicing of XBP1 was done using 8-formyl-7-hydroxy-4-methylcoumarin (4µ8c) or 6-Bromo-2-hydroxy-3-methoxybenzaldehyde (Sigma-Aldrich). Both inhibitors inhibit IRE1-mediated splicing of XBP1 in other cell types. For inhibition in NK-cells, 60µM was added to cultures at the same time as target cells.

Dicer-substrate short interfering RNA (DsiRNA) by Integrated DNA Technologies (IDT) was also used to knockdown IRE1α (official gene symbol: ERN1) mRNA at three different locations (Exons 4, 10 and 22). A negative control (NC) DsiRNA was used to demonstrate the specific effect of DsiRNA targeting IRE1α. HEK293T cells were treated with IRE1α DsiRNA or NC DsiRNA (24pmol) using RNAiMax and IRE1α mRNA quantified after 24 hours. DsiRNA treated HEK293T cells were stimulated with TG to assess IRE1α activation by qRT-PCR of XBP1s mRNA and miR-34a-5p. To independently validate the association between IRE1α, XBP1 and miR-34a-5p, HEK293T cells were treated with thapsigargin (TG, 100nM) and IRE1α inhibitor (4µ8c, 60µM) for 2 hours and *XBP1s* mRNA and miR-34a-5p expression quantified.

#### Dual luciferase reporter assay for miRNA target validation

A dual luciferase reporter system was used to validate the direct interaction of miR-34a-5p and the predicted target site on the 3’UTR of PD-1. Dual-Luciferase® Reporter Assay System with psiCHECK2 (Ψ) vector was used to assess the changes in the expression of chimeric luciferase constructs mediated by miRNA oligonucleotide mimics in HEK293T cells. PsiCHECK2 (Ψ) vector encodes two reporter genes: (1) the firefly (*Photinus pyralis*) luciferase (Fluc), and (2) *Renilla reniformis* luciferase (Rluc) that contains multiple cloning sites for insertion of 3’UTR or miRNA regulating elements (MRE) from target predictions. Fluc expression is used as an internal control for the plasmid transfection, whereas the chimeric Rluc expression (Rluc-3’UTR or -MREs) is affected by specific miRNA binding.

miRNA mimics were designed by IDT. Two different plasmid reporter constructs were generated: a construct carrying small synthetic sequences of putative target site plus flanking restriction sites and a positive-control plasmid carrying a perfect complementary sequence of mature miRNA. Plasmid reporter construction was performed as previously outlined (4, 5), plasmid DNA and miRNA mimics were co- transfected into HEK293T cells grown in DMEM with 10% fetal bovine serum (FBS) using Lipofectamine3000 (ThermoFisher) transfection reagent. The ratios of expression levels of chimeric Rluc and Fluc (used as transfection normalizer) were compared between miR-34a-5p mimics and negative control mimics (cel-miR-67-5p) for the dual luciferase assay. miRNA mimics rescue assay was performed using a miR-34a inhibitor (anti-miR-34a-5p) and a negative control inhibitor (anti-miR-NC). Measurements were performed 24 hours post-transfection. Dual-Luciferase® Reporter Assay System by Promega was used to measure the chimeric Rluc and Fluc expression levels and the luminescence measurements were acquired in Biotek Synergy 2 microplate reader.

#### Luciferase reporter assay for validation of XBP1 binding sites in PD-1 gene

pGL3 basic vector luciferase reporter (Promega) was used to assess the changes in the expression of chimeric luciferase constructs transfected to HEK293T cells using Lipofectamine3000. pGL3 basic vector encodes the reporter firefly (*Photinus pyralis*) luciferase (Fluc) gene that contains multiple cloning sites in its promoter for insertion of regulatory regions from target predictions. pGL3 basic vector constructs carrying amplified genomic (upstream, intronic and downstream) regions and small synthetic sequences of putative XBP1s target site plus flanking restriction sites. Plasmid reporter construction was performed as previously outlined (4, 5), plasmid DNA was transfected into HEK293T cells grown in DMEM with 10% fetal bovine serum (FBS) using Lipofectamine3000 (ThermoFisher) transfection reagent. The ratio of expression levels of chimeric Fluc was compared between TG (XBP1s induction) and DMSO treatments. Measurements were performed 24 hours post-transfection and 6hours post TG treatment. Dual-Luciferase® Reporter Assay System by Promega was adapted to measure the Fluc expression and the luminescence was acquired in Biotek Synergy 2 microplate reader.

#### miRNA mimic treatment in NK-cells

PD-1 expressing NK-cell leukaemia cell-line KHYG-1 grown in RPMI 1640 (Gibco Life Technologies) with 10% FBS (Gibco Life Technologies) and 100 U/mL interleukin-2 was treated with miRNA mimics. Knockdown of target mRNA and protein was measured by qRT-PCR and flow cytometry respectively. For delivery of miRNA mimics into KHYG-1 cells we used Lipofectamine RNAiMAX Transfection Reagent as per manufacturer's protocol. Target mRNA and protein knockdown were quantified by comparing KHYG-1 cells treated with miR-34a-5p relative to a non-specific control (cel-miR-67-5p). For qRT-PCR, we used miRNeasy Kit (Qiagen) to isolate the target mRNA, cDNA was synthesised with Qiagen miScript PCR system and the relative delta Ct was calculated by normalising to GAPDH.

#### Biotinylated miRNA – mRNA pulldown transcriptome analysis

Biotinylated microRNA-mRNA pulldown experiments were used to identify bound mRNA transcripts of miR-34a-5p. Briefly, biotinylated miRNAs (miR-34a-5p and Cel-67-5p) were transfected into PD-1 expressing KHYG-1 NK cell-line and captured along with their targeted transcripts to magnetic streptavidin beads (Invitrogen Dynabeads M-280).

The pulldown protocol was primarily adapted from Wani and Cloonan (5), and Cristino and Barchuk (4). The samples were reverse transfected in a 75 $cm^2$ flask $4x10^6$ cells with biotinylated miRNA with lipofectamine 3000 (Thermo Fisher) and grown for 24 h before pulldown. Four biological replicates, from four independent transfection and pulldown experiments, were made for each miRNA (miR-34a-5p and cel-67-5p). Pre-pulldown samples (one fourth of the whole samples) were saved for normalisation control. RNA was purified (both pre-pulldown and pulldown samples) with RNAeasy columns (QiAGEN) using their RNA clean up protocol, reverse transcribed to cDNA using QuantiTect Reverse Transcription Kit and predicted miR-34a-5p targets (XBP1 and PD-1) and non-target (SDHA) mRNAs were quantified using qRT-PCR in ViiA™ 7 Real-Time PCR System (Thermo Fisher Scientific). Relative delta Ct was calculated by normalising to respective pre-pulldown samples and later calibrated biotinylated miR-34a-5p treatment against biotinylated cel-67-5p treatment.

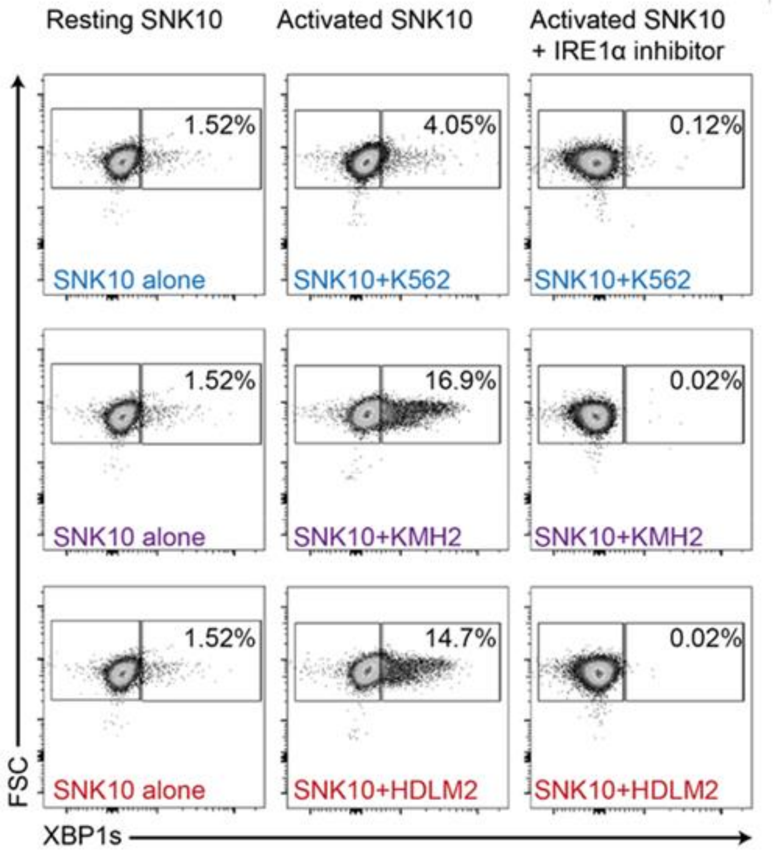


**Figure S1: IRE1α pathway activation in NK-cells stimulated with target cells.** Representative FACS plots showing XBP1s protein expressed in SNK10 cells following activation with target cells (K562, KMH2 and HDLM2), with or without IRE1α inhibitor (4μ8c 60μM)

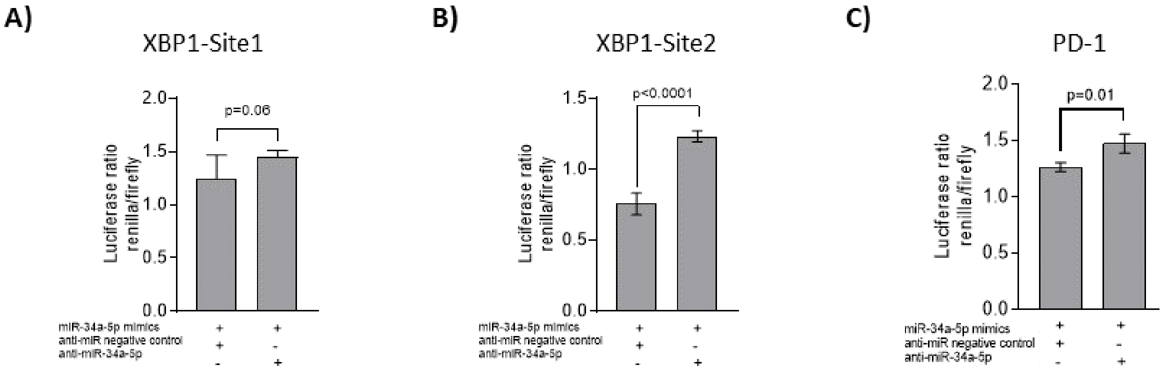


**Figure S2: Rescue assay showing blockade of miR-34a-5p mimics repression over XBP1 and PD-1 target sites.** Dual-luciferase reporter assay showing rescue effect of miR-34a-5p inhibitor (anti-miR-34a-5p) over miR-34a-5p target sites in XBP1 Site 1 **(A)** and Site 2 **(B)**. HEK293T cells treated with miR-34a-5p and anti-miR-34a-5p reversed the reduction in luciferase signal caused by miR-34a-5p mimics (plus negative control, anti-miR NC). **(C)** Dual-luciferase reporter assay of miR-34a-5p inhibitor (anti-miR-34a-5p) showing rescue effect over miR-34a-5p target sites in PD-1.

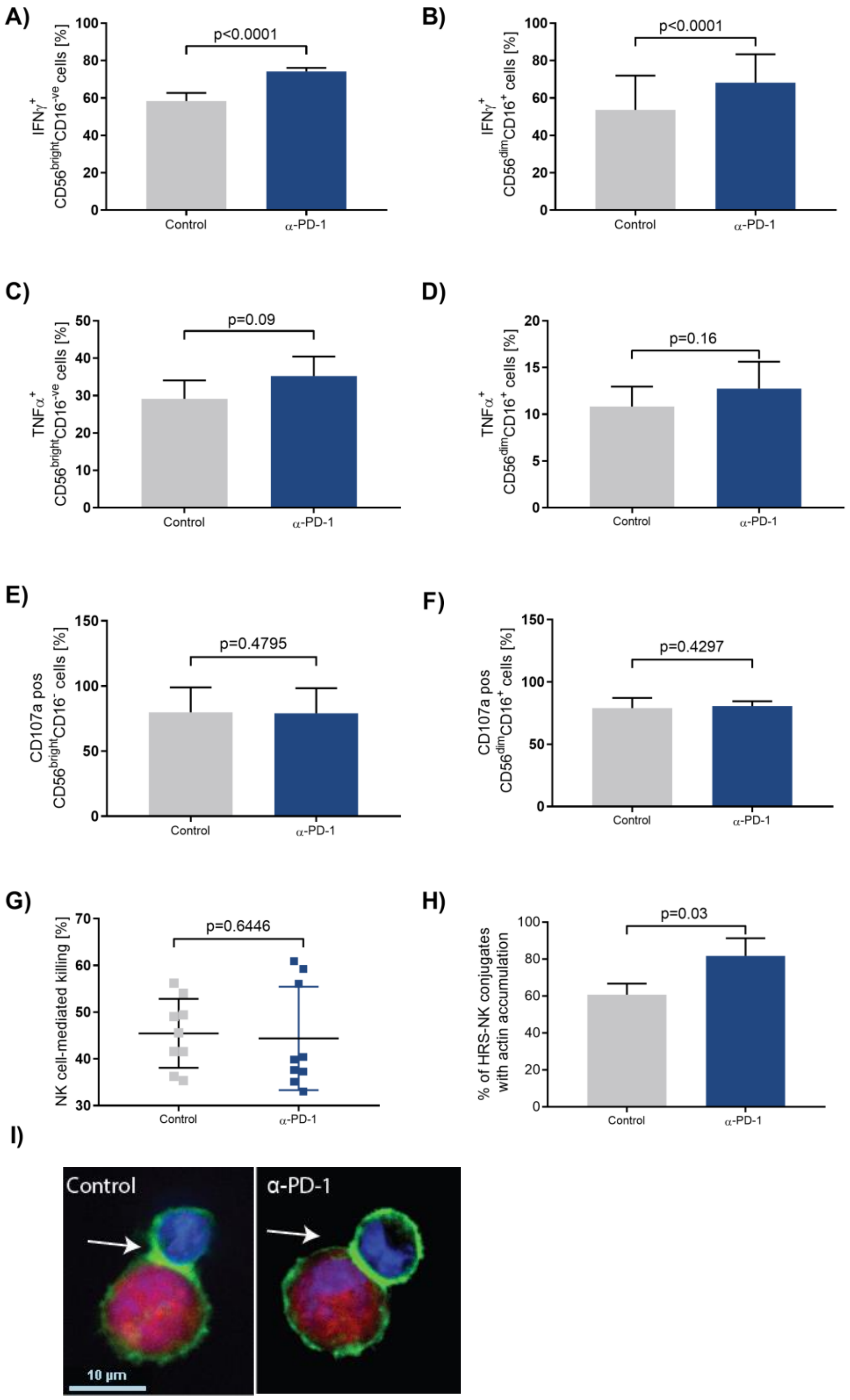

A)
IFNγ+ CD56brightCD16-ve cells [%]
p<0.0001
Control
α-PD-1
B)
IFNγ+ CD56dimCD16+ cells [%]
p<0.0001
Control
α-PD-1
C)
TNFα+ CD56brightCD16-ve cells [%]
p=0.09
Control
α-PD-1
D)
TNFα+ CD56dimCD16+ cells [%]
p=0.16
Control
α-PD-1
E)
CD107a pos CD56brightCD16- cells [%]
p=0.4795
Control
α-PD-1
F)
CD107a pos CD56dimCD16+ cells [%]
p=0.4297
Control
α-PD-1
G)
NK cell-mediated killing [%]
p=0.6446
Control
α-PD-1
H)
% of HRS-NK conjugates with actin accumulation
p=0.03
Control
α-PD-1
I)
Control
α-PD-1
10 µm

**Figure S3: The effector functions of NK-cells from patients with HL upon α-PD-1 treatment.** After stimulation with HDLM2 cells for 6 hours. cytokines (**A,B**) IFNγ and (**C,D**) TNFα were assessed by FACS in CD56brightCD16-ve and CD56dimCD16+ cells. PD-1 blockade significantly enhanced IFNγ release in both CD56brightCD16-ve and CD56dimCD16+ subsets but α-PD-1 treatment in pNK-cells did not enhance TNFα release in pNK-cell subsets compared to Control (DMSO and IgG4 isotype). **(E,F)** PD-1 blockade did not enhance NK-cell de-granulation in **(E)** $CD56^{bright}CD16^{ve-}$ or (**F**) $CD56^{dim}CD16^{+}$subset of NK-cells. **(G)** No significant difference in cytotoxicity was observed upon PD-1 blockade in both NK-cell subset. **(H)** NK-cell immune synapse formation between NK-cell and HRS cells is restored after PD-1 blockade with increased F-actin polymerization at the NKIS. **(I)** Representative images of NK-cell immune synapse formation between NK-cell (DAPI, blue) and HRS cells (APC, red) showing the expanded NK-cells from patients with HL stimulated with HDLM2 for 6 hours in the presence of PD-1 blockade (α-PD-1). Arrows indicate actin accumulation (green) at the NK-cell/HRS synapse site. In the control assays, HRS target cells and pNK-cells were co-incubated with immunoglobulin (Ig) G4 isotype control. Original magnification ×63.

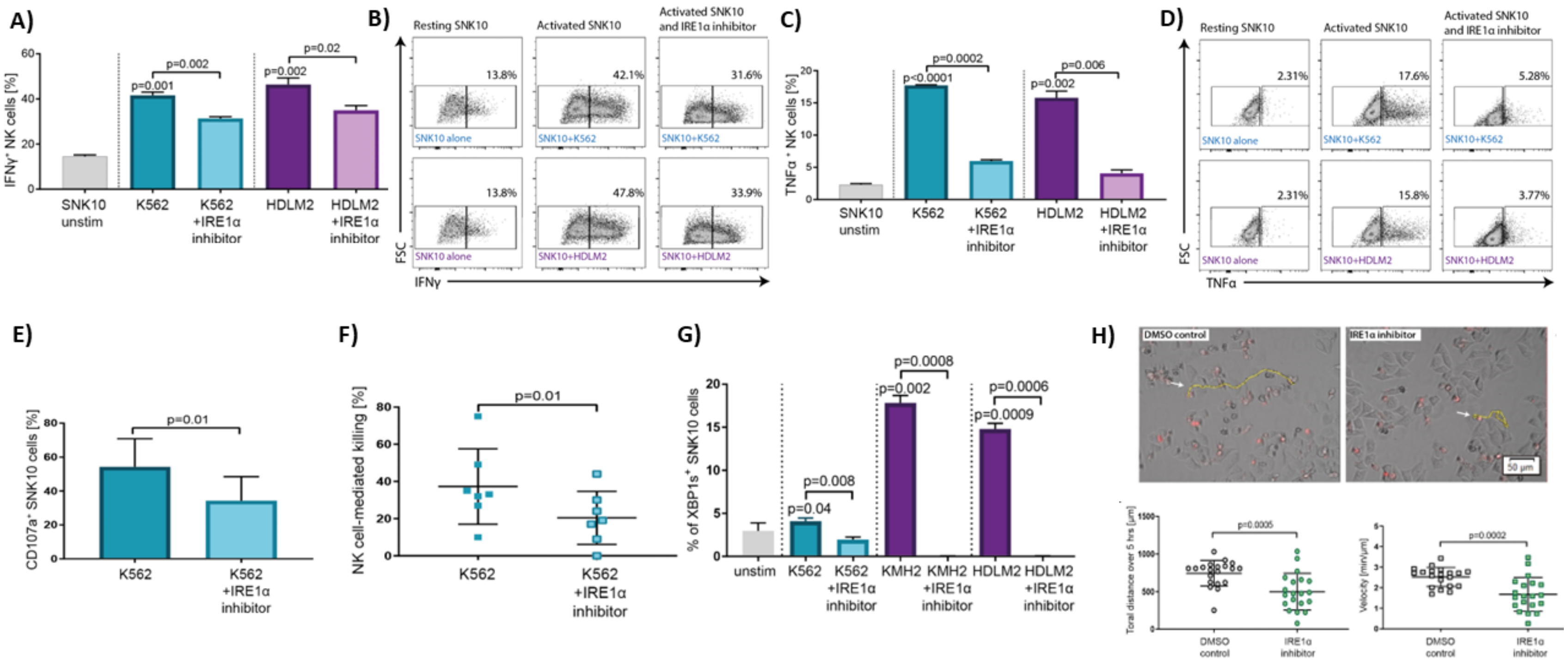
A)
IFNγ⁺ NK cells [%]
p=0.002
p=0.001
p=0.02
p=0.002
SNK10 unstim
K562
K562 +IRE1α inhibitor
HDLM2
HDLM2 +IRE1α inhibitor
B)
Resting SNK10
Activated SNK10
Activated SNK10 and IRE1α inhibitor
13.8%
42.1%
31.6%
SNK10 alone
SNK10+K562
SNK10+K562
13.8%
47.8%
33.9%
SNK10 alone
SNK10+HDLM2
SNK10+HDLM2
FSC
IFNγ
C)
TNFα⁺ NK cells [%]
p=0.0002
p<0.0001
p=0.006
p=0.002
SNK10 unstim
K562
K562 +IRE1α inhibitor
HDLM2
HDLM2 +IRE1α inhibitor
D)
Resting SNK10
Activated SNK10
Activated SNK10 and IRE1α inhibitor
2.31%
17.6%
5.28%
SNK10 alone
SNK10+K562
SNK10+K562
2.31%
15.8%
3.77%
SNK10 alone
SNK10+HDLM2
SNK10+HDLM2
FSC
TNFα
E)
CD107a⁺ SNK10 cells [%]
p=0.01
K562
K562 +IRE1α inhibitor
F)
NK cell-mediated killing [%]
p=0.01
K562
K562 +IRE1α inhibitor
G)
% of XBP1s⁺ SNK10 cells
p=0.0008
p=0.002
p=0.0006
p=0.0009
p=0.008
p=0.04
unstim
K562
K562 +IRE1α inhibitor
KMH2
KMH2 +IRE1α inhibitor
HDLM2
HDLM2 +IRE1α inhibitor
H)
DMSO control
IRE1α inhibitor
50 μm
Toral distance over 5 hrs [μm]
p=0.0005
DMSO control
IRE1α inhibitor
Velocity [min/μm]
p=0.0002
DMSO control
IRE1α inhibitor

**Figure S4: IRE1α blockade impairs several hallmark NK-cell immune effector functions.** (**A**) Intracellular IFNγ was quantified by FACS in SNK10 cell-line treated with DMSO or IRE1α inhibitor (60µM of 4µ8c) in response to K562 and HDLM2 cells. A significant reduction in IFNγ release was observed upon IRE1α-blockade in the SNK10 cell-line stimulated with K562 and HDLM2 target cells. P values above a single bar significance when compared to unstimulated control. **(B)** Intracellular representative FACS plots showing IFNγ in resting and activated SNK10 cells with and without IRE1α inhibitor. (**C**) TNFα protein expression significantly reduced upon IRE1α-blockade in the SNK10 cell-line stimulated with K562 and HDLM2 target cells**.** P values above a single bar significance when compared to unstimulated control. **(D)** Intracellular representative FACS plots showing TNFα in resting and activated SNK10 cells with and without IRE1α inhibitor. **(E)** A significant reduction in CD107a protein expression was observed in activated SNK10 cells treated with IRE1α inhibitor suggesting that IRE1α inhibitor reduces the ability of SNK10 cell-line to de-granulate when exposed to K562 target cell. (**F**) IRE1α inhibitor significantly reduces the ability of SNK10 cell-line to lyse K562 target cells. (**G**) Inhibition of IRE1α with 4µ8c downregulates XBP1s protein in NK-cells stimulated with different target cells. P values above a single bar significance when compared to unstimulated control. **(H)** Representative single-cell tracking maps of SNK10 cells trajectories are shown (yellow line), with total distance and mean velocity. Each data point represents 1 single randomly selected NK-cell tracked over 5 hours. NK-cell motility of untreated SNK10 cells (in red) and IRE1α inhibited SNK10 cells (4µ8c 60µM) towards HeLa target cells was assessed using live cell imaging and time-lapse microscopy. Results of total distance and velocity are displayed as the mean ± SD of at least 6 representative regions of interest (ROIs) in time-lapse videos obtained from 3 independent experiments. As expected, SNK10 cells treated with IRE1α inhibitor showed significant reduction in total distance and velocity compared to DMSO control treatment.

A)

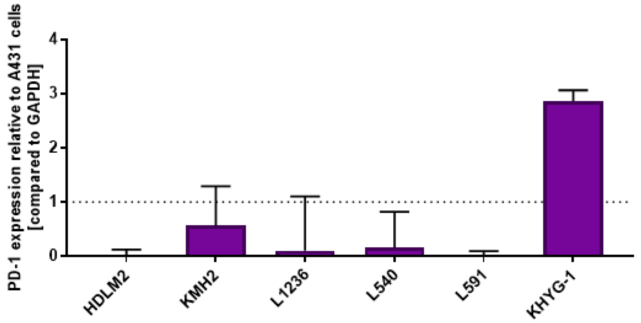


B)

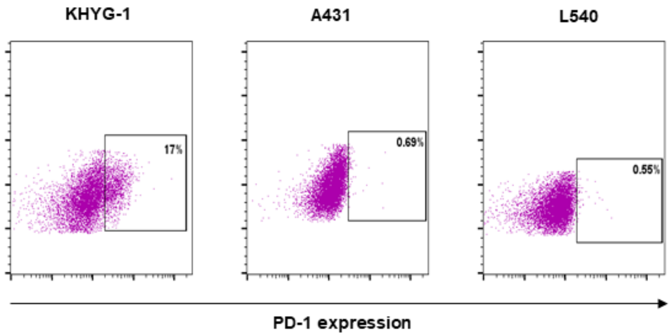


C)

| Cell line | HDLM2 | KMH2 | L540 | L591 | L1236 |
|---|---|---|---|---|---|
| PD-1 expression (mRNA) | - | - | - | - | - |
| PD-1 expression (protein) | - | - | - | - | - |
| Type | Nodular sclerosis | Mixed cellularity progressing to lymphocyte depletion | Nodular sclerosis | Nodular sclerosis | Mixed cellularity |
| Stage | IV | IV | IVB | IVB | IV |
| EBV status | - | - | - | + | - |
| Immunology | CD30 + | CD30 + | CD30 + | CD30 + | CD30 - |

D)

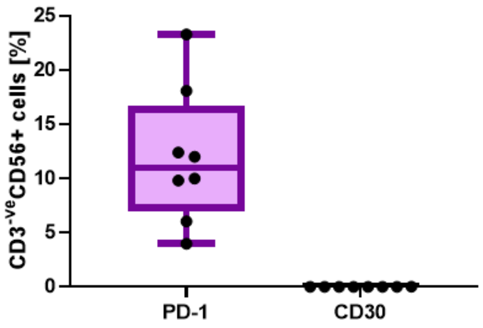

**Figure S5: Expression of PD-1 and CD30 in HL cell-lines and pNK-cells in patients with HL showing trogocytosis is not occurring between HL cell-lines and pNK-cells. (A)** PD-1 mRNA expression was tested in a range of HL cell-lines. None of the tested cell-lines expressed PD-1 mRNA relative to the negative control A431 cell-line. KHYG-1 was used as a positive control. **(B)** Representative flow cytometry plots of PD-1 protein expression in KHYG-1 (positive control), A431 (negative control) and L540 HL cell-line. **(C)** Summary of important characteristic details of tested HL cell-lines including PD-1 mRNA and protein expression, as well as HRS cells marker (CD30) expression. **(D)** *Ex-vivo* expression of PD-1 and CD30 in pNK-cells in pre-therapy blood from patients with HL. pNK-cells from patients with HL expressed PD-1 but not the HRS cells marker CD30 indicating that PD-1 expression on pNK-cells is not due to trogocytosis.

**Table S1: Computational predictions of miR-34a-5p target sites in PD-1 and XBP1 3'UTR using miRanda.**

Putative miRNA target sites (free energy < -18kcal/mol and score >= 140) were predicted in the 3'UTR of PDCD1-201 (PD-1) and XBP1-201 (sites overlap both XBP1u and XBP1s isoforms) transcripts described in GENCODE database (GRCh37 release 38). Only miR-34a-5p target sites were found in the 3'UTR of these genes.

| Target transcript | Free energy | Score | Target site (5'→ 3') | Target start position | Target stop Position |
|---|---|---|---|---|---|
| **PDCD1-201** | -20.48 | 147 | AGGGCCAGATGCAGTCACTGCTT | 1172 | 1194 |
| **XBP1-201** | -20.53 | 148 | GCTTTCATCCAGCCACTGCCC | 1056 | 1076 |
| **XBP1-201** | -18.84 | 154 | TTCCTTGACTATTACACTGCCT | 1256 | 1277 |
